# Proteomic characterization of the *Mycobacterium marinum*-containing vacuole in *Dictyostelium discoideum*

**DOI:** 10.1101/592717

**Authors:** Aurélie Guého, Cristina Bosmani, Jahn Nitschke, Thierry Soldati

## Abstract

*Mycobacterium tuberculosis*, the causative agent of tuberculosis, is able to manipulate the phagosome compartment in which it resides in order to establish a permissive replicative compartment called the *Mycobacterium*-containing vacuole (MCV). *Mycobacterium marinum*, a fish pathogen and a close relative of the tuberculosis group, is able to infect the free-living amoeba and professional phagocyte *Dictyostelium discoideum* and to manipulate its phagosome maturation. By using this host-pathogen model system, we have established an innovative process to isolate MCVs. This procedure allowed us to isolate the *M. marinum*-MCV at 1, 3 and 6 hours post infection to study the early *M. marinum*-MCV proteome. By using isobaric labelling and mass spectrometry, we quantitatively compared the proteomic composition of those MCVs isolated at different stages of the early infection phase to understand how *M. marinum* impacts on this compartment to divert it from the normal phagosomal pathway. Furthermore, we also compared the manipulated compartment *M. marinum*-MCV to non- or less manipulated compartments containing different mycobacteria strains: the non-pathogenic *M. smegmatis*, the avirulent *M. marinum*-L1D or the attenuated *M. marinum*-RD1.

## INTRODUCTION

*Mycobacterium tuberculosis* (Mtb) is the causative agent of tuberculosis, an infectious disease still responsible of 1.5 million deaths in 2020 (Geneva: World Health Organization, 2021). Once taken up by macrophages, it has the particularity to be able to manipulate the phagosomal compartment where it resides to prevent its killing and degradation, and consequently establish a successful infection (Russell, 2007). *Mycobacterium marinum* is closely related to Mtb and can cause a similar disease in cold blooded animals (Tobin and Ramakrishnan, 2008).

*Dictyostelium discoideum* is a professional phagocyte that uses phagocytosis to feed on bacteria. Molecular components of its phagosomal pathway are highly conserved with those of human phagocytic cells (Boulais et al., 2010; Dunn et al., 2018). Thanks to its ease-of-use and the numerous genetic tools existing, it has been widely used to dissect the phagosome maturation process. It has also emerged as a macrophage model organism to decipher the infection fate of phagocytosed or intraphagosomal pathogens (Hagedorn and Soldati, 2007; Lampe et al., 2016; Solomon et al., 2000).

The well-established *D. discoideum*-*M. marinum* system has allowed the deciphering of different key steps of a mycobacterial infection (Hagedorn and Soldati, 2007). Indeed, like Mtb, *M. marinum* can manipulate the maturation of the phagosome where it resides and tailor it into a replication-permissive compartment, the mycobacteria-containing vacuole or MCV. It modifies the phosphoinositide composition of its containing-vacuole (Koliwer-Brandl et al., 2019). It can prevent the acidification of the compartment where it resides by inhibiting the recruitment of the H^+^-vATPase or by promoting its recycling from the MCV (Koliwer-Brandl et al., 2019; Kolonko et al., 2014). It can induce the relocalisation of the host lipid droplets to the close vicinity of its MCV in order to use their content (Barisch et al., 2015; Barisch and Soldati, 2017). At later times of infection, post-lysosomal makers such as the copper transporter p80 (Kolonko et al., 2014) or vacuolin, the functional homolog of flotillin (Hagedorn, 2007) accumulate on the MCV. However, the lysosomal enzyme CatD does not (Cardenal-Muñoz et al., 2017; Hagedorn and Soldati, 2007). H^+^-vATPase retrieval from the MCV and no delivery of lysosomal enzymes allow the mycobacteria to reside in a non-acidic and non-degradative post-lysosome-like compartment where it can replicate. Eventually, *M. marinum* induces the rupture of its MCV to escape into the cytosol where it can access nutrients (Hagedorn and Soldati, 2007), and then disseminate to neighbouring cells through ejectosomes (Hagedorn et al., 2009) or cell lysis (López-Jiménez et al., 2019). This escape in the cytosol occurs at later time of infection, but *M. marinum* starts damaging its MCV membrane very early during the infection process. To prevent this escape, both the ESCRT and the autophagy machineries of the host cell participate in the sealing of the wounded membrane (Cardenal-Muñoz et al., 2017; López-Jiménez et al., 2018).

Several virulence factors, necessary for the establishment of a successful infection, have been identified in *M. marinum*. One of them is the *mag* 24-1 gene which encodes a PE-PGRS protein: a protein with the characteristic Pro-Glu (PE) motif in N-term and high polymorphic GC-rich sequences (PGRS). It is necessary for the intracellular replication of *M. marinum* in human and frog macrophages and in *D. discoideum*. The *M. marinum*-L1D mutant, whose *mag* 24-1 gene is inactivated by transposon insertion, has lost this ability (Hagedorn and Soldati, 2007; Ramakrishnan et al., 2000). Another important pathogeny locus in the *M. marinum* genome is the Region of Difference 1 (RD1) locus. It encodes virulence factors and the Type VII secretion system (TSS7) ESX-1 necessary for their secretion into the MCV lumen. One of these factors is EsxA, a 6 kDa peptide with membranolytic properties. The RD1 locus is then essential for MCV membrane damage. However, those damages are not only necessary for MCV rupture at later time points of infection (Hagedorn et al., 2009), but also during the very early phase of infection for the establishment of the replicative niche (Cardenal-Muñoz et al., 2017; López-Jiménez et al., 2018; Tan et al., 2006). Indeed, the *M. marinum*-RD1 mutant can survive in its MCV but is not able to fully arrest phagosome maturation (Tan et al., 2006), to recruit lipid droplets around its MCV (Barisch et al., 2015). It cannot induce damage nor escape from its MCV (Cardenal-Muñoz et al., 2017; López-Jiménez et al., 2018), nor replicate nor form ejectosomes to disseminate to neighbouring cells (Hagedorn et al., 2009).

Despite numerous studies, it is still not clear if the *M. marinum*-MCV remains an early phagosome or if it has a more complex identity. Furthermore, it is still poorly understood how the virulent mycobacteria manipulate the phagosome maturation program. To answer these questions, we decided to study exhaustively and quantitatively the proteomic composition of the *M. marinum*-MCV in *D. discoideum* and its changes during the first six hours of infection, which we term the “manipulation phase”. We also compared the proteomic composition of this manipulated compartment to the proteomic composition of MCVs containing a non-pathogenic mycobacterium (*M. smegmatis*), an avirulent *M. marinum* (L1D), or an attenuated *M. marinum* (*M. marinum*-RD1).

## RESULTS AND DISCUSSION

### Establishment of a new procedure to isolate mycobacteria-containing vacuoles

In order to decipher the MCV proteome, the first step of this study has been to establish an experimental strategy to isolate pure fractions of those compartments from infected *D. discoideum* cells. We decided to adapt the efficient method to isolate pure fractions of latex beads-containing phagosomes (LPCBs) described previously (Dieckmann et al., 2012). This method is based on the light buoyant density and flotation properties of latex beads. As a brief reminder of that generic protocol, *D. discoideum* cells are fed with latex beads, phagosome maturation is stopped at different stages and cells are mechanically broken to liberate the LBCPs, which are recovered by flotation on a sucrose gradient. Our adapted strategy (**Figure 1A**), is to float up MCVs on a sucrose gradient with the help of latex beads. For this, we established conditions to adsorb latex beads onto mycobacteria by hydrophobic interactions with the waxy surface of the mycobacteria. The optimal conditions are based on a standard protocol used to adsorb proteins on latex beads. After several washing steps of both latex beads and mycobacteria in borate buffer, beads and mycobacteria were mixed and incubated on a rotating wheel. After 2 h, we could observe the formation of bacteria and beads complexes (BBCs) (**Figure 1B, and S1B**).

**Figure 1.**
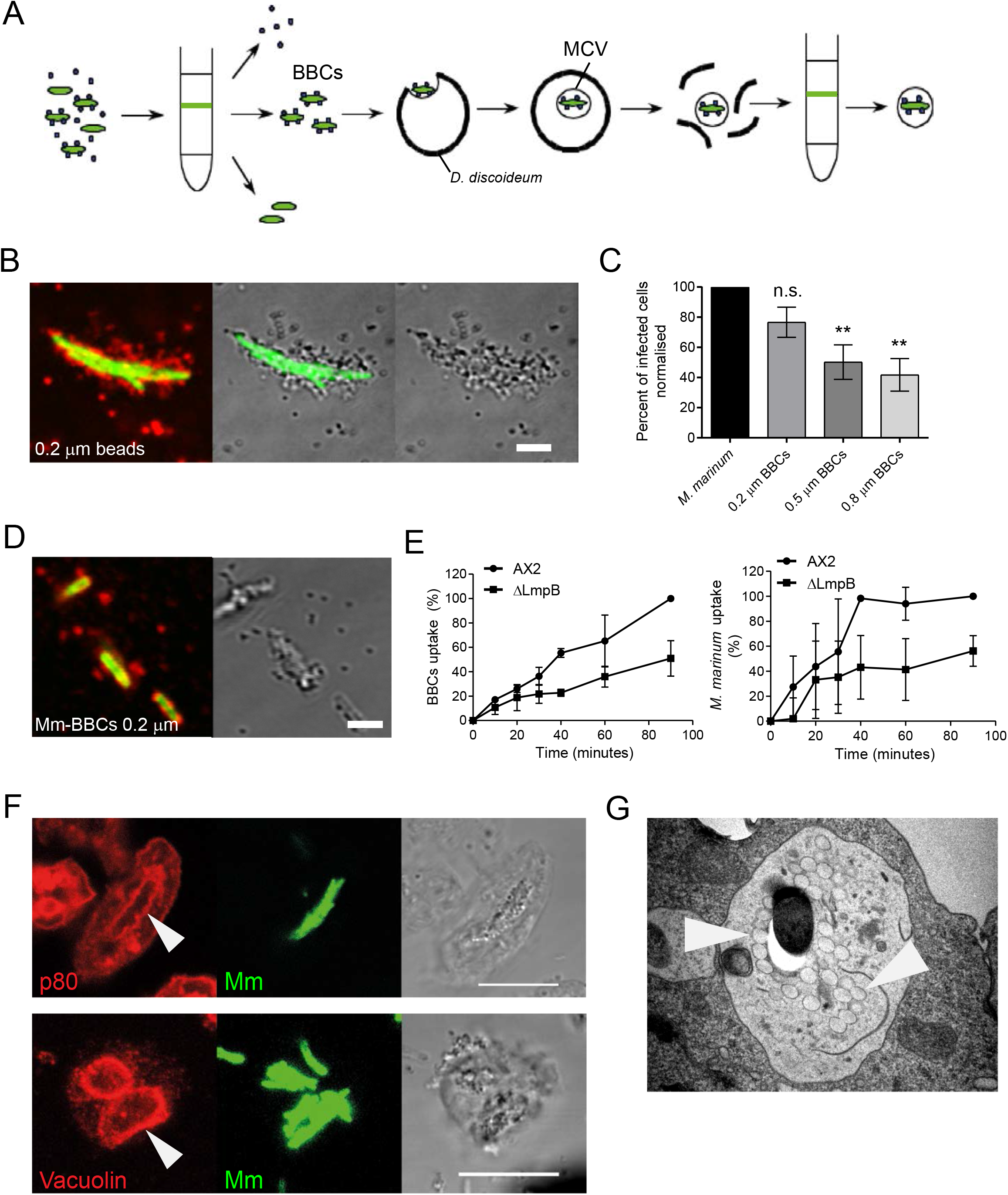
BBCs are recognized and phagocytosed as mycobacteria by *D. discoideum*. **A**. Experimental strategy used to isolate mycobacteria-containing vacuoles (MCVs) from infected cells. Latex beads were adsorbed onto mycobacteria. A pure inoculum of bacteria and bead complexes (BBCs) was prepared and used to infect *D. discoideum* cells. At times of interest, cells were homogenized, and the homogenate was loaded onto a sucrose gradient and ultracentrifuged. Compartments containing BBCs float at the top of the sucrose gradient and were recovered. **B**. *M. marinum*-GFP-BBCs prepared with 0.2 μm latex beads. For ease of visualisation, TRITC-coupled antibodies (red) were adsorbed onto latex beads during BBCs preparation. Scale bar, 1 μm. **C**. Wild type cells were infected with *M. marinum*-GFP or *M. marinum*-GFP-BBCs prepared with different sizes of latex beads. The proportion of infected cells was determined by FACS. The values were normalized to the number of infected cells obtained for the infection performed with non-coated *M. marinum*-GFP. Bars represent mean and SD of 4 independent experiments (one-way ANOVA, **p<0.01). **D**. BBCs were prepared with *M. marinum-GFP* and 0.2 μm latex beads and stained for anti-mycobacterial cell wall material with a cocktail of antibodies (anti-*M. leprae* membrane antigens, anti-*M. leprae* surface antigens, anti-*M. leprae* cell wall associated antigens, red). Scale bars, 2 μm. **E**. The uptake of non-coated *M. marinum*-GFP (right) and of 0.2 μm-*M. marinum*-GFP-BBCs (left) by wild type and *lmpB-ko* cells was measured by FACS. Relative fluorescence was normalized to the value obtained for wild type cells at 90 min. The curve represents the mean and SD of 3 independent experiments. **F**. Wild type cells were infected with 0.2 μm-*M. marinum*-GFP-BBCs. At 21 hpi, cells were immunostained against p80 or vacuolin. Arrowheads indicate p80- or vacuolin-positive membranes around *M. marinum*-BBCs. Scale bar, 10 μm. **G**. Wild type cells were infected with 0.2 μm-*M. marinum*-GFP-BBCs. At 21 hpi, cells were processed for Transmission Electron Microscopy (TEM). Arrowheads indicate 0.2 μm latex beads.

Different bead sizes (0.8 μm, 0.5 μm and 0.2 μm), and different bead:bacteria ratios were tested. The ratios were adapted according to the bead size: 2 and 15 for 0.8 μm beads, 30 and 90 for 0.5 μm beads, and 100 and 500 for 0.2 μm beads (**Figure S1A**). Interestingly, the highest ratios had a “declumping” effect on the mycobacteria, as they prevented mycobacteria from sticking to each other, and allowed the formation of BBCs containing single bacteria. The BBCs obtained with 0.8 μm beads were large, about twice the size of a single mycobacteria, while 0.5 μm and 0.2 μm beads did not increase drastically the size of mycobacteria.

In order to completely cover mycobacteria with beads, a large excess of latex beads has to be used, resulting in a profusion of free beads. To isolate fractions of pure MCVs without contamination of phagosomes containing the free beads, un-adsorbed beads have to be removed from the BBC preparation before infecting cells. To achieve this, a one-step sucrose gradient was sufficient (**Figure S1C**). After ultracentrifugation, BBCs could be collected at the 20% −60% sucrose interphase, whereas free beads floated up at the 20% sucrose-medium interphase.

To confirm the possibility to use BBCs to isolate MCVs, the flotation property of BBCs on the standard sucrose gradients used to isolate LBCPs was monitored (**Figure S1D**). Most of the BBCs concentrated at the 10%-25% sucrose interphase, like the LBCPs. Only few BBCs, probably with a lower ratio of beads and too dense to float to the top of the gradient, were observed at the other interphases. To increase the yield of MCV isolation, the sucrose gradient used for LBCPs (10%, 25%, 35%, 60%) was adjusted to 10%, 30%, 40%, 60%. These pilot experiments allowed us to prepare BBC inoculates devoid of free beads and with the property to float up on sucrose gradients similar to those used for LBCPs isolation.

### BBCs are recognized and phagocytosed like “normal” mycobacteria

BBCs are slightly bigger than single mycobacteria and have an irregular shape. In order to test whether *D. discoideum* could phagocytose them, cells were infected with non-coated *M. marinum*-GFP or *M. marinum*-GFP BBCs prepared with three different sizes of beads. At 0.5 hpi, the percentage of infected cells was measured by flow cytometry (**Figure 1C**). All three sizes of BBCs were efficiently ingested, however, the percentage of infected cells was bead-size-dependent. The highest percentage of internalized BBCs was obtained with 0.2 μm beads. Therefore, we decided to use these BBCs for further experiments.

Mycobacteria in BBCs are totally covered with latex beads. To test whether BBCs are recognized by *D. discoideum* as latex beads or as mycobacteria, immunofluorescence was performed with a cocktail of antibodies against components of the mycobacterial cell wall. Interestingly, beads covering mycobacteria and free beads incubated with mycobacteria were detected by this cocktail (**Figure 1D**). This indicates that some components of the cell wall diffuse and cover the adsorbed beads. In addition, materials from the mycobacteria cell wall seem to be shed in the incubation buffer and adsorbed onto free beads. To confirm that the *M. marinum*-BBCs are recognized by *D. discoideum* as mycobacteria, and not inert beads, internalization of BBCs was monitored in *ImpB*-ko cells, which are specifically defective for the uptake of mycobacteria and not latex beads (Sattler et al., 2018). As for noncoated *M. marinum, M. marinum*-BBCs ingestion was significantly impaired in *ImpB*-ko cells (**Figure 1E**). These results suggest that BBCs are recognized as mycobacteria.

Several hours after uptake by *D. discoideum*, *M. marinum*-BBCs were observed in closed compartments (**Figure1G**), that were positive for p80 and vacuolin (**Figure 1F**), known markers of the MCV (Hagedorn and Soldati 2007). Latex beads were also observed inside these compartments, indicating that mycobacteria and latex beads remain in the same compartment even several hours postinfection. Overall, our results show that BBCs are recognized and taken up like free mycobacteria and reside in compartments similar to MCVs.

### Mycobacteria in the BBCs are still alive and infectious

During preparation of BBCs, mycobacteria are subjected to different mechanical, osmotic and chemical stresses. To test whether this has an impact on the infectivity of mycobacteria, *D. discoideum* cells were infected with different non-coated GFP-expressing mycobacteria or BBCs prepared with different strains: the virulent *M. marinum*, the avirulent *M. marinum*-L1D and the non-pathogenic *M. smegmatis*. These strains are known to follow different fates and their infection cycles have been previously described (Hagedorn and Soldati, 2007). The infection course of each bacterium or BBC was then monitored by flow cytometry (**Figure 2A, B**). As previously described, the non-pathogenic *M. smegmatis* was rapidly killed, and the population of infected cells progressively decreased until almost no infected cells were detected at 18 hours postinfection (hpi). The population of cells infected with the avirulent *M. marinum*-L1D also decreased at 6 hpi, while the population of extracellular mycobacteria increased. *M. marinum*-L1D is known to be exocytosed within few hours and re-ingested before being killed, resulting in a strong decrease of the population of infected cells and concomitant increase of extracellular mycobacteria at 18 hpi (Hagedorn and Soldati 2007). On the contrary, the virulent *M. marinum* is able to establish a successful infection, with a high proportion of infected cells remaining at 21 hpi. The infections performed with BBCs showed similar behaviour. *D. discoideum* cells infected with *M. smegmatis*-BBCs or *M. marinum*-L1D-BBCs managed to cure the infection in several hours, whereas cells remained infected 21 hpi with *M. marinum*-BBCs. This indicated that the fate of the different BBCs is dependent on the used mycobacteria strain and that BBCs follow the same infection course as the mycobacteria they contain. These data also corroborate our previous results (**Figures 1F-G and 2D**) showing that *M. marinum*-BBCs remained in a compartment as late as 21 hpi, in contrast to inert particles, which are exocytosed in 1 or 2 hours (Hagedorn and Soldati 2007).

**Figure 2.**
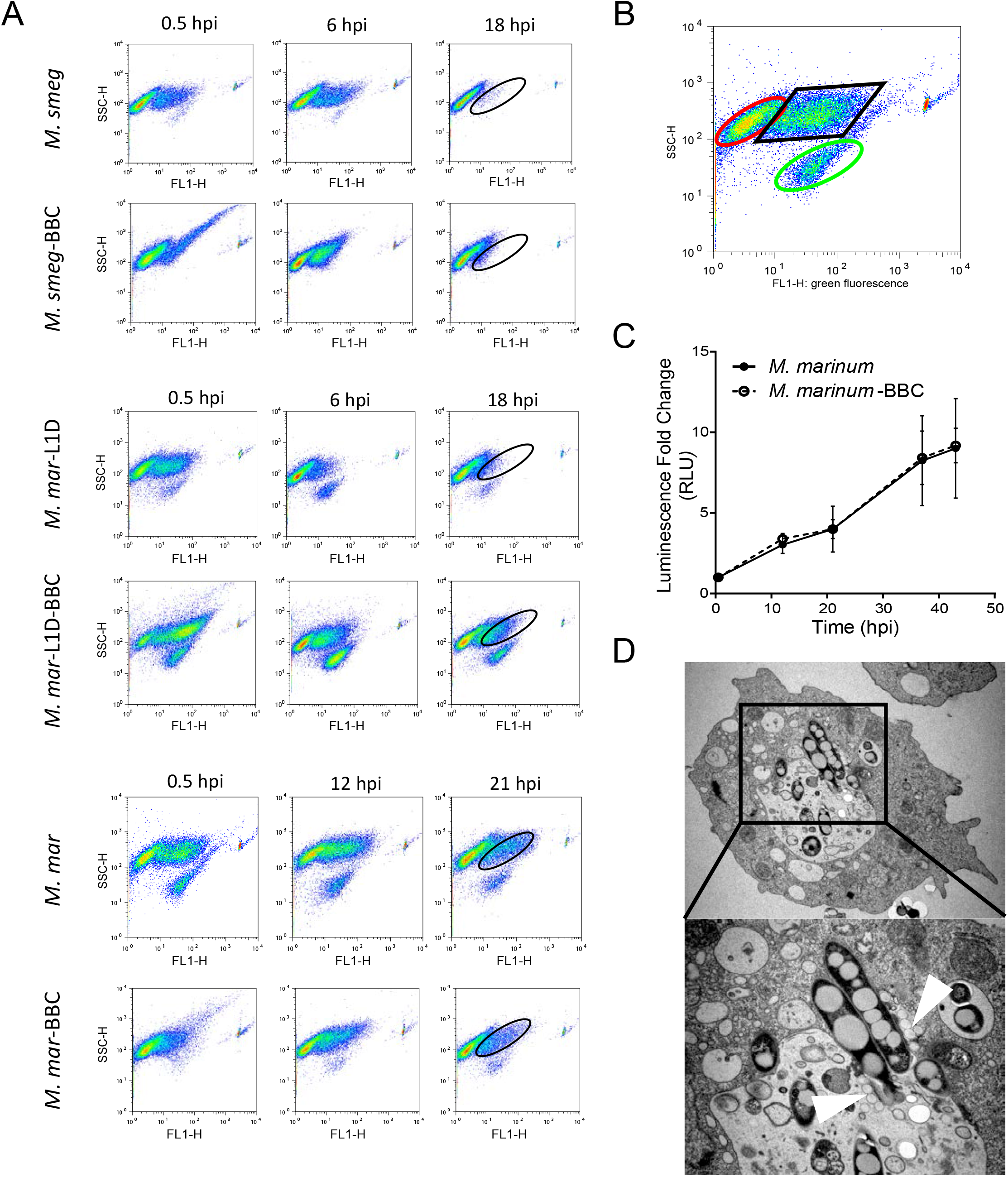
BBCs are infectious and behave as noncoated mycobacteria. **A**. Wild type cells were infected with *M. smegmatis*-GFP, *M. marinum*-L1D-GFP, *M. marinum*-GFP or BBCs prepared with these different strains. After removing uningested bacteria or BBCs, infection was monitored by FACS. Representative SSC (side scatter) vs FL-1 (GFP fluorescence) plots are presented. **B**. Three different populations can be distinguished: non-infected cells (red), infected cells (black) and extracellular mycobacteria or BBCs (green). **C**. Wild type cells were infected with *M. marinum*-lux. Intracellular mycobacteria growth was monitored by measuring the luminescence at indicated times. The curves represent the mean fold increase of luminescence and SD of 3 independent experiments. **D**. Wild type cells were infected with 0.2 μm-*M. marinum*-GFP-BBCs. At 21 hpi, cells were processed for TEM. Arrowheads indicate 0.2 μm latex beads.

To test whether mycobacteria in the BBCs were still metabolically active and able to grow, *D. discoideum* cells were infected with BBCs prepared with luminescent *M. marinum* or non-coated *M. marinum*-lux. Growth was then followed by measuring the luminescence emitted by *M. marinum*-lux (**Figure 2C**). No intracellular growth difference was observed between *M. marinum* from BBCs and non-coated *M. marinum*, demonstrating that mycobacteria are alive in the BBCs.

It has been shown that at later stages of infection, *M. marinum* breaks its compartment to escape to the cytosol (Hagedorn and Soldati, 2007; López-Jiménez et al., 2018). By electron microscopy, virulent *M. marinum* from BBCs were observed escaping their compartment at 21 hpi, indicating that the beads do not disturb the normal fate of *M. marinum* nor its virulence, as it is still able to damage the phagosome membrane (**Figure 2D**).

These results indicate that despite the stresses incurred by the mycobacteria during the preparation of BBCs, mycobacteria incorporated in the BBCs are still alive, infectious and virulent. Furthermore, the beads do not disturb the steps of a normal infection cycle.

### Preliminary characterization of isolated MCVs

We performed an initial characterization of MCVs after isolation at 1 hpi, as this was the earliest time point accessible technically. The recovered interphase contained a high number of BBCs (**Figure 3A**). Since uningested BBCs were washed away after the spinoculation, this recovered material was likely entirely BBCs-containing compartments. Very little other cellular material than BBCs was observed, indicating that the recovered BBC-containing MCVs were not significantly contaminated by other cellular organelles.

**Figure 3.**
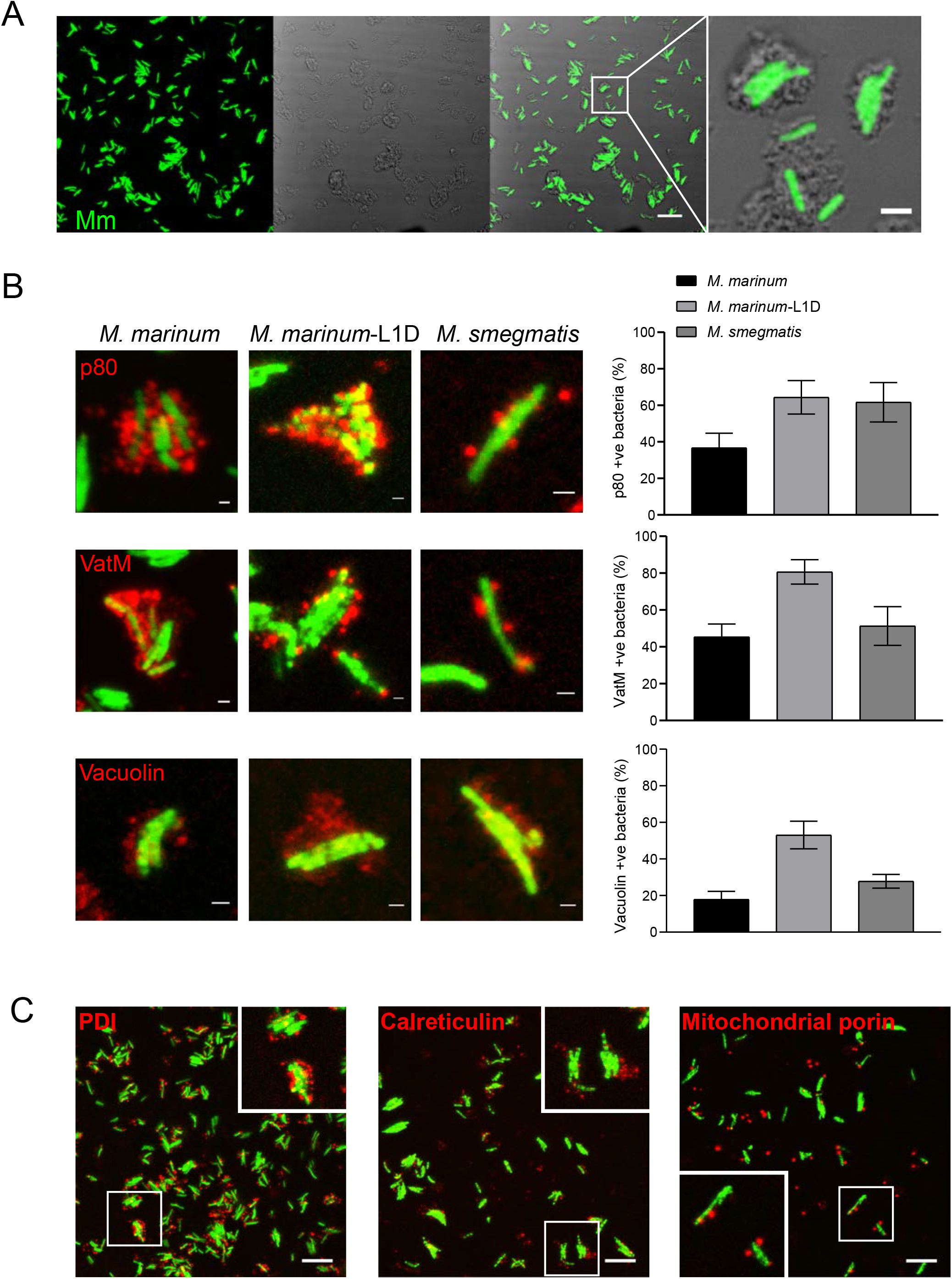
Vacuoles containing different mycobacteria strains can be isolated. **A**. Wild type cells were infected with 0.2 μm-*M. marinum-GFP-* BBCs. At 1 hpi, cells were homogenized. After incubation with ATP, the cell homogenate was ultracentrifuged on a sucrose gradient. The fraction recovered at the 10 %-30 % interphase was observed by microscopy. Scale bar, 10 μm; in the zoom-in, 2 μm. **B**. MCVs containing 0.2 μm-BBCs prepared with *M. marinum*-GFP, *M. marinum*-L1D or *M. smegmatis* were isolated from wild type cells and immunostained against p80, VatM or vacuolin. The number of compartments positive for the different markers were counted for each mycobacteria strain. Scale bars, 1 μm. Graphs represent mean and SEM of 2 independent experiments. **C**. MCVs containing 0.2 μm-*M. marinum*-GFP-BBCs were isolated from wild type cells and immunostained against PDI, calreticulin or mitochondrial porin (red). Scale bars, 10 μm.

MCVs containing BBCs generated with two other mycobacteria, *M. marinum*-L1D or *M. smegmatis*, were also isolated at 1 hpi. Immunostainings were performed on the recovered material to test the presence of known markers of the MCV around BBCs (**Figure 3B**). This indicated the presence of a phagosomal membrane around the recovered BBCs and confirmed that MCVs can be isolated with this new procedure. Furthermore, the immunofluorescence revealed some differences between the manipulated MCV of *M. marinum*, and the non-manipulated MCVs of *M. marinum*-L1D and *M. smegmatis* at 1 hpi. Indeed, only few *M. marinum* MCVs were positive for the two post lysosome markers p80 and vacuolin (36% and 18% respectively), whereas about 60% of the *M. marinum*-L1D and *M. smegmatis* MCVs were positive for p80, and 53% of *M. marinum*-L1D MCVs and 27% of *M. smegmatis* MCVs were positive for vacuolin. Interestingly, 45% of the *M. marinum* MCVs were positive for the H^+^-vATPase, confirming the observations of a previous study that showed that more than 50% of *M. marinum* MCVs transiently acquire the H^+^-vATPase during the first hour of infection (Hagedorn and Soldati 2007). However, higher percentages of MCVs positive for H^+^-vATPase were found for *M. marinum*-L1D and *M. smegmatis*, 80% and 51% respectively. Altogether, these results indicate that already at 1 hpi, *M. marinum* has manipulated the compartment where it resides to divert it from the normal phagosome maturation process. In addition, *M. marinum* appears to reside in a compartment with some of the characteristics of an early phagosome. On the contrary, at 1 hpi, the avirulent *M. marinum*-L1D and the non-pathogenic *M. smegmatis* are in MCVs with characteristics of mature phagosomes. These results also demonstrate that, despite the beads and the potential stresses incurred by the mycobacteria during the different steps of the procedure, the identity of the isolated MCVs depends on the mycobacteria strain.

To verify the purity of the recovered MCVs, *M. marinum*-BBCs were isolated at 1hpi. The presence of markers for the endoplasmic reticulum (ER, monitored for protein disulfide-isomerase, PDI, and calreticulin) and for mitochondria (mitochondrial porin) was then evaluated by immunofluorescence (**Figure 3C**). Note that PDI and calreticulin are also detected transiently at early phagosomes (Dieckmann et al., 2012). Interestingly, a high proportion of *M. marinum*-BBCs were positive for both ER markers at 1hpi, especially PDI, confirming the early phagosome identity of the compartment where it resides. ER fragments not associated with BBCs were only rarely detected. Few BBCs were positive for mitochondrial porin. These results indicate that the new procedure established here allows the isolation of relatively pure fractions of MCVs. Overall, we demonstrated that BBCs are an appropriate and efficient tool to further characterize the MCV proteome.

### *M. marinum* MCV proteome

Biological duplicates of MCVs containing *M. marinum*-BBCs where isolated at early times of infection: 1, 3 and 6 hpi. A comparative quantitative proteomic strategy using tandem mass tag (TMT) technology (sixplex TMT® MS/MS) was used to study their composition. 1606 *D. discoideum* proteins (**Table S1**) and 563 *M. marinum* proteins were identified and constitute the early MCV proteome. Among the *D. discoideum* proteins of the early MCV, 758 were also found in the phagosome proteome (Dieckmann et al., 2012) and 875 in the macropinosome proteome (Journet et al., 2012), whereas 525 of these proteins were found to be unique to the early MCV proteome (**Figure 4A**). The high proportion of proteins in common between these different compartments indicates that the MCV, despite being manipulated by virulent mycobacteria, remains a compartment of phagosomal/endosomal origin.

**Figure 4.**
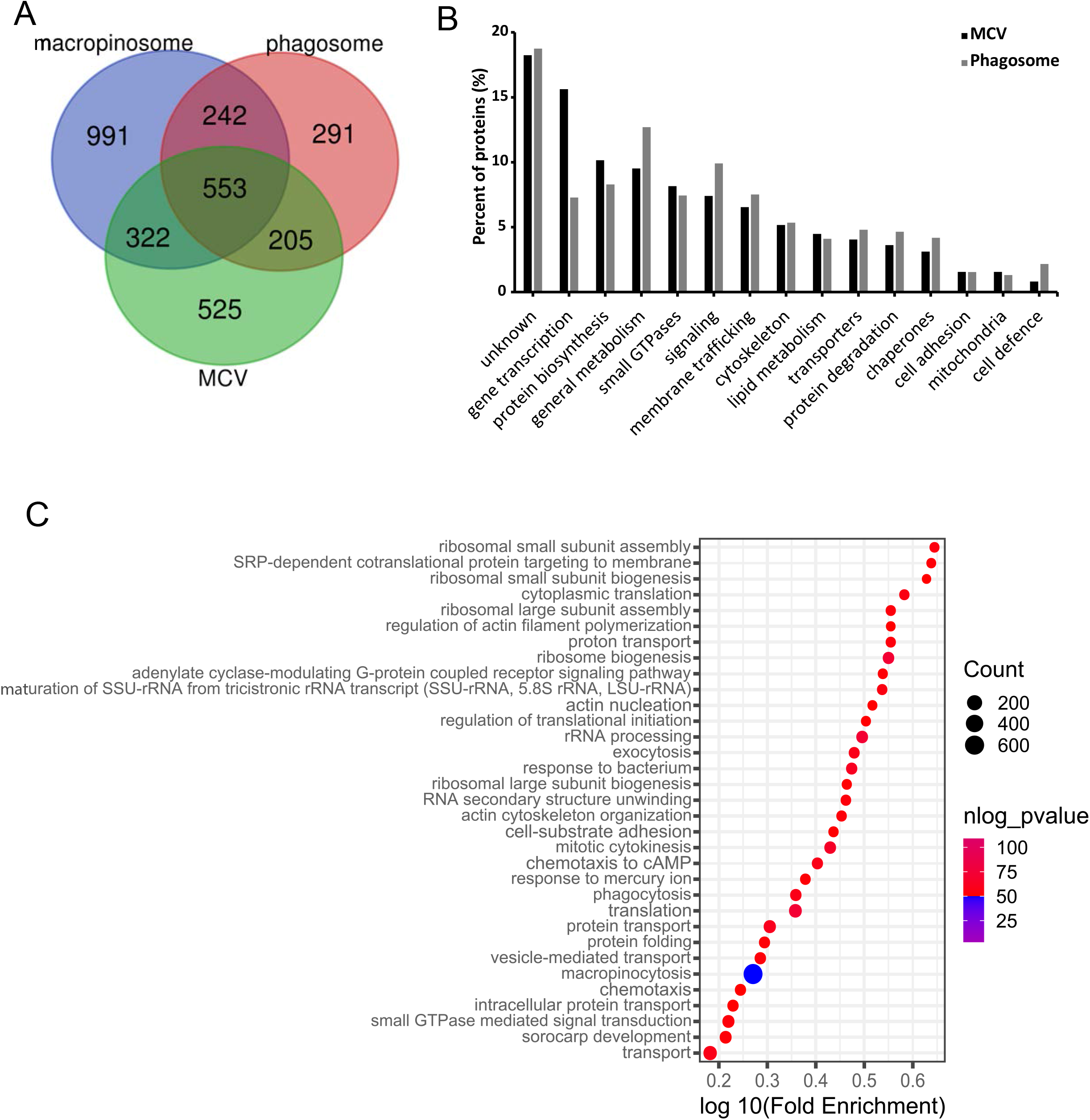
Proteomic analysis of the *M. marinum*-MCV during the early phase of infection (1 to 6 hpi). *M. marinum*-MCV were isolated at 1, 3 and 6 hpi. After labelling with TMT, equal amounts of each sample were mixed. This pool was analysed by LC-MS/MS. **A**. Venn diagram depicting the number of proteins commonly found in the *M. marinum*-MCV proteome and in the published phagosome proteome, and macropinosome proteome. **B.** Functional classification of the 1606 *D. discoideum* proteins identified in the *M. marinum*-MCV proteome by mass spectrometry. As a comparison, results from a previous phagosome proteome analysis are also presented, and classified using the same classification terms. Percentages of identified proteins in the different classes are indicated. **C.** Functional clustering of the *M. marinum*-MCV proteome with the help of the DAVID functional annotation clustering tool. The default parameters of the program were used and the categories Gene Ontology “biological process” and KEGG were selected. Enriched GO-terms are presented according to their enrichment score and p-value (p-value<10^-3^). Size of the bubble represents the number of proteins in the cluster.

To identify pathways involved in the establishment of a permissive niche, proteins identified in the early MCV were manually sorted in different classes according to their function (**Figure 4B**). They have also been functionally annotated and clustered with the help of the DAVID functional annotation clustering tool (**Figure 4C** and **Table S2**) (Huang et al., 2009; Sherman et al., 2022). Comparison of the MCV proteome functional classification with the previously published phagosome proteome classification (Dieckmann et al., 2012) indicated that overall, these compartments are very similar (**Figure 4B**), confirming again that the MCV conserves some of its phagosomal identity. In accordance with this, the most significantly enriched GO terms with the highest number of proteins were “macropinocytosis”, “phagocytosis” and “response to bacterium”.

However, some differences were clearly apparent between the phagosome and the MCV proteomes. The two highest represented classes in the MCV were “gene transcription” and “protein biosynthesis”. Both were more represented in the MCV proteome than in the phagosome proteome. These classes of proteins are generally delivered through the autophagic pathway to phagosomes for degradation. Their high representation in the MCV proteome could indicate an accumulation of those proteins in a non-degradative phagosome, and thus reflect the manipulation of this compartment by *M. marinum*. This was confirmed by the functional annotation analysis, where “translation”, “ribosome biogenesis” and “rRNA processing” were among the most significantly enriched categories for GO terms, and “ribosome” and “RNA transport” for KEGG pathways. Also related to translation machinery, the GO terms “maturation of SSU-rRNA”, “RNA secondary structure unwinding”, “cytoplasmic translation”, “ribosomal small subunit assembly”, “ribosomal small subunit biogenesis”, “regulation of translational initiation” and “ribosomal large subunit biogenesis” were found significantly enriched in the MCV (**Figures 4B-C**).

Two other classes were found slightly more represented in the MCV than in the phagosome proteome: “small GTPases” and “lipid metabolism”. “Small GTPase mediated signal transduction” was also found significantly enriched among the functionally annotated GO terms of the *M. marinum*-MCV proteome.

Interestingly, classes of proteins involved in “general metabolism”, “signalling”, “membrane trafficking”, “cytoskeleton”, “transporters”, “protein degradation”, “chaperones” and “cell defence” were less represented in the MCV than in the phagosome. All these classes are involved in or reflect the maturation of the phagosome (signalling, cytoskeleton and membrane trafficking) and in the establishment of an acidic and degradative compartment (transporters, protein degradation, cell defence). This indicates again that the MCV is a phagosome with a less degradative lumenal environment. Despite an under-representation in the MCV compared to the phagosome, these classes were also found significantly enriched in the functional annotation clustering analysis of the MCV proteome: the cluster “transport” and “protein transport”, GO terms “proton transport”, “vesicle-mediated transport”, “intracellular protein transport”, “adenylate cyclase-modulations G-protein coupled receptor signalling pathway”, “actin cytoskeleton organization”, “actin nucleation”, “regulation of actin filament polymerization” and “exocytosis” and KEGG pathways “oxidative phosphorylation” and “phagosome”.

### Early *M. marinum*-MCV maturation

Two different ratios were calculated for each of the 1606 proteins of the early *M. marinum*-MCV proteome: 3 hpi/1 hpi and 6 hpi/1 hpi. For each comparison, a ratio significance cut-off threshold was calculated (**Table S5)** (Tiberti et al., 2010). From this, 139 proteins were found differentially abundant in the *M. marinum*-MCV between 1 hpi and 3 hpi, and 146 proteins between 1 and 6 hpi. (**Table S3**). The log2 ratios of those differentially abundant proteins were plotted (**Figure 5A**). Log2 ratios that did not pass the cut-off threshold were noted as 0. From this graph, six different kinetic profiles are distinguishable during the first 6 hours of MCV establishment, compared to the composition at 1 hpi: proteins continuously depleted, proteins continuously accumulating, proteins transiently depleted at 3 hpi, proteins transiently accumulating at 3 hpi, proteins starting to accumulate only from 3 hpi onward, and finally, proteins starting to be depleted only from 3 hpi onward. 133 proteins were depleted from the MCV between 1 and 3 hpi, and 132 between 3 and 6 hpi, whereas only 6 proteins accumulated in the compartment between 1 and 3 hpi, and 59 between 3 and 6 hpi, indicating that the MCV is established mainly by depleting proteins compared to its composition one hour after formation. In order to identify the pathways involved in the establishment of a permissive MCV, the proteins accumulating or depleted from the compartment at the different time points were manually sorted and functionally annotated and clustered using the DAVID functional annotation clustering tool (**Figure 5B, C, D, E)** (Huang et al., 2009; Sherman et al., 2022). Overall, all the protein classes identified in the early MCV proteome were affected during the first 6 hours of infection (**Figure 5B**).

**Figure 5.**
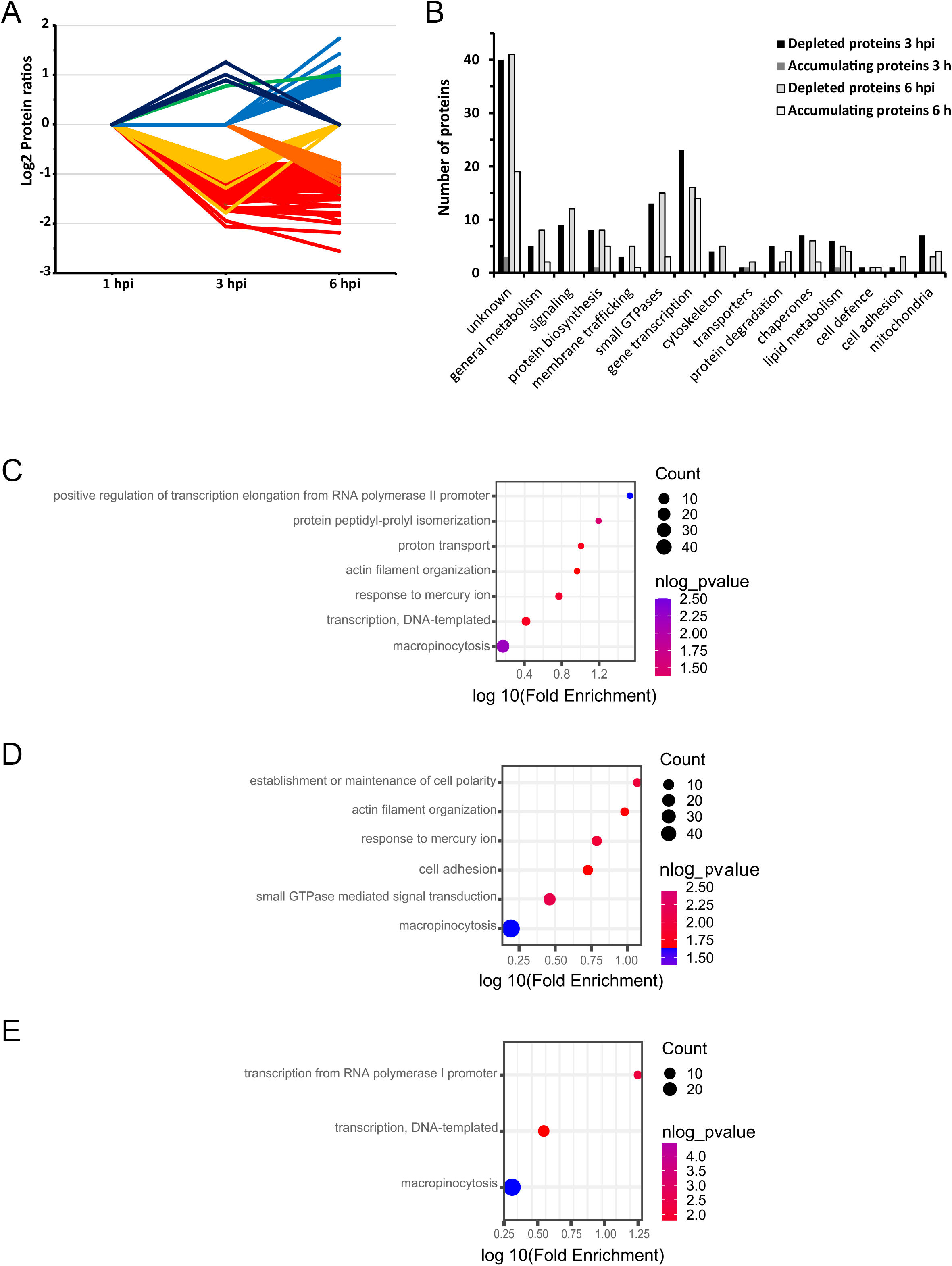
Temporal modification of the *M. marinum*-MCV during the early phase of infection (1 to 6 hpi). *M. marinum*-MCV were isolated at 1, 3 hpi and 6 hpi. After labelling with TMT, equal amounts of each sample were mixed. This pool was analysed and quantified by LC-MS/MS. Both *M. marinum*-MCV isolated at 3 and 6 hpi were compared to *M. marinum*-MCV isolated at 1 hpi. For each identified protein, the protein ratios of the corresponding comparison were calculated. **A.** Log2 ratios of the 232 *D. discoideum* MCV proteins significantly differentially abundant during the early phase of infection. Log2 ratios no significantly different from 0 were noted as 0. **B.** MCV proteins significantly differentially abundant during the early phase of infection were functionally classified using the same classification terms as in Figure 4. *M. marinum*-MCV proteins significantly depleted at 3 hpi **(C)** at 6 hpi **(D)** or accumulating at 6 hpi **(E)** were functionally annotated using the DAVID functional annotation clustering tool. The default parameters of the program were used and the category Gene Ontology “biological process” was selected. Enriched GO-terms are presented according to their enrichment score and p-value. Size of the bubble represents the number of proteins in the cluster.

Protein classes the most depleted at 3 and 6 hpi were very similar. The “Gene transcription” class was the most represented. The cluster “transcription” was also among the most significantly enriched GO-terms among the proteins depleted at 3 hpi. Proteins of this class being mainly delivered to phagosomes for degradation via the autophagic pathway, this might indicate that the MCV stops the normal phagosome maturation by preventing the fusion with any other intracellular compartments. Some other classes like “protein biosynthesis” and “chaperones” and GO term like “protein peptidyl-prolyl isomerization” have similar fate and were also found significantly enriched among the depleted proteins.

The classes “signalling”, “membrane trafficking” and “small GTPases” were strongly depleted from the MCV. Consistent with this, “small GTPase mediated signal transduction” was among the most significantly enriched GO terms depleted at 6 hpi. These classes and GO terms are all involved in phagosome maturation. Their depletion from the early MCV again indicates that the maturation of this compartment and its fusion with other endosomal compartments is arrested. However, not only the fusion is blocked. Indeed, proteins involved in fission were also found in those classes (Dynamin A, retromer subunits Vps52, Vps53). Their depletion might prevent the removal of early phagosomal markers, a necessary step for further MCV maturation, potentially explaining how the MCV identity is reported to resemble the early phagosome stage.

The class “cytoskeleton” and the corresponding significantly enriched GO term “actin filament organization” were also depleted from the MCV both at 3 and 6 hpi. An actin coat is indeed retained around the MCV during the first hours of infection and necessary to prevent fusion with lysosomes. This coat is then lost at later times of infection (Kolonko et al., 2014). These classes and GO terms might reflect the progressive removal of the actin coat.

The GO term “proton transport” was significantly enriched among the depleted proteins at 3 hpi. The depletion of components of the H^+^ v-ATPase allows to avoid or to rapidly stop the acidification of the MCV lumen. The class “general metabolism” was depleted indicating that the entire cell metabolism could be affected during mycobacterial infection. More precisely, the “lipid metabolism” class was depleted at 3 and 6 hpi. Among those proteins were oxysterol binding proteins 8 and 12, which are involved in sterol transport. Their depletion could prevent removal of lipids from the MCV. Furthermore, from 3 hpi, this class accumulated again in the MCV. However, instead of lipid transporter, proteins really involved in lipid metabolism accumulated from 3 hpi. This would allow the mycobacteria to access and use host lipids (Barisch and Soldati 2017).

As the “lipid metabolism” class, classes “protein biosynthesis” and “gene transcription” were depleted from 1 to 3 hpi, and then among the most accumulating ones from 3 to 6 hpi. Consistent with this, the cluster “transcription” was also significantly enriched at 6 hpi. This indicates that after preventing fusion with endo-lysosomal compartments, the MCV starts again to interact with intracellular compartments after 3 hpi. However, the accumulation of those classes of proteins indicates that the lumen of the MCV remains non-degradative and that it fuses only with non-degradative compartments. The accumulation, to a lesser extent, of the classes “membrane trafficking” and “small GTPases” from 3 hpi confirmed this hypothesis.

It is interesting to notice that there was only one protein continuously accumulating on the MCV during the 6 first hours of infection. This protein was a phox-domain containing protein that appears to be a homolog of human Snx4. Through its association with the protein KIBRA, it is involved in the sorting of the transferrin receptor from early endosomes to the endocytic recycling compartment (Solis et al., 2013). Interestingly, the *D. discoideum* KIBRA homolog, a WW domain-containing protein, was also found differentially abundant during MCV establishment. However, it was continuously depleted, potentially blocking Snx4 function and further phagosome maturation, and consequently leading to the accumulation of Snx4 on the MCV.

In conclusion, two phases could be distinguished during the first six hours of MCV establishment. From 1 to 3 hpi, fusion of the MCV with endo-lysosomal compartments is blocked. Proteins involved in phagosome maturation, and in establishment of a degradative environment are removed. Fission is also blocked, preventing removal of early phagosomal markers. From 3 to 6 hpi, fusions with non-degradative intracellular compartments might be increased again. As lipid metabolism proteins start accumulating, these fusions could allow the delivery of lipids or any nutrient necessary for the growth of the mycobacteria.

### *M. marinum*-MCV comparison with “non-manipulated”-MCVs during the early maturation phase

Biological duplicates of MCVs containing *M. marinum*, *M. smegmatis* or *M. marinum*-RD1 were isolated at 1 and 6 hpi. MCVs containing *M. marinum*-L1D were isolated only at 1 hpi since *M. marinum*-L1D is rapidly exocytosed (Hagedorn and Soldati 2007). A comparative quantitative proteomic strategy was used to compare *M. marinum*-MCV composition to other *Mycobacteria*-MCVs at 1 and at 6 hpi. For each comparison, protein ratios were calculated. A significance ratio cut-off threshold was calculated (**Table S5**) in order to select proteins significantly enriched or depleted on the *M. marinum*-MCV (**Table S4**). For each comparison, the number of proteins significantly enriched or depleted on the *M. marinum*-MCV is presented in **Figure 6A**.

**Figure 6.**
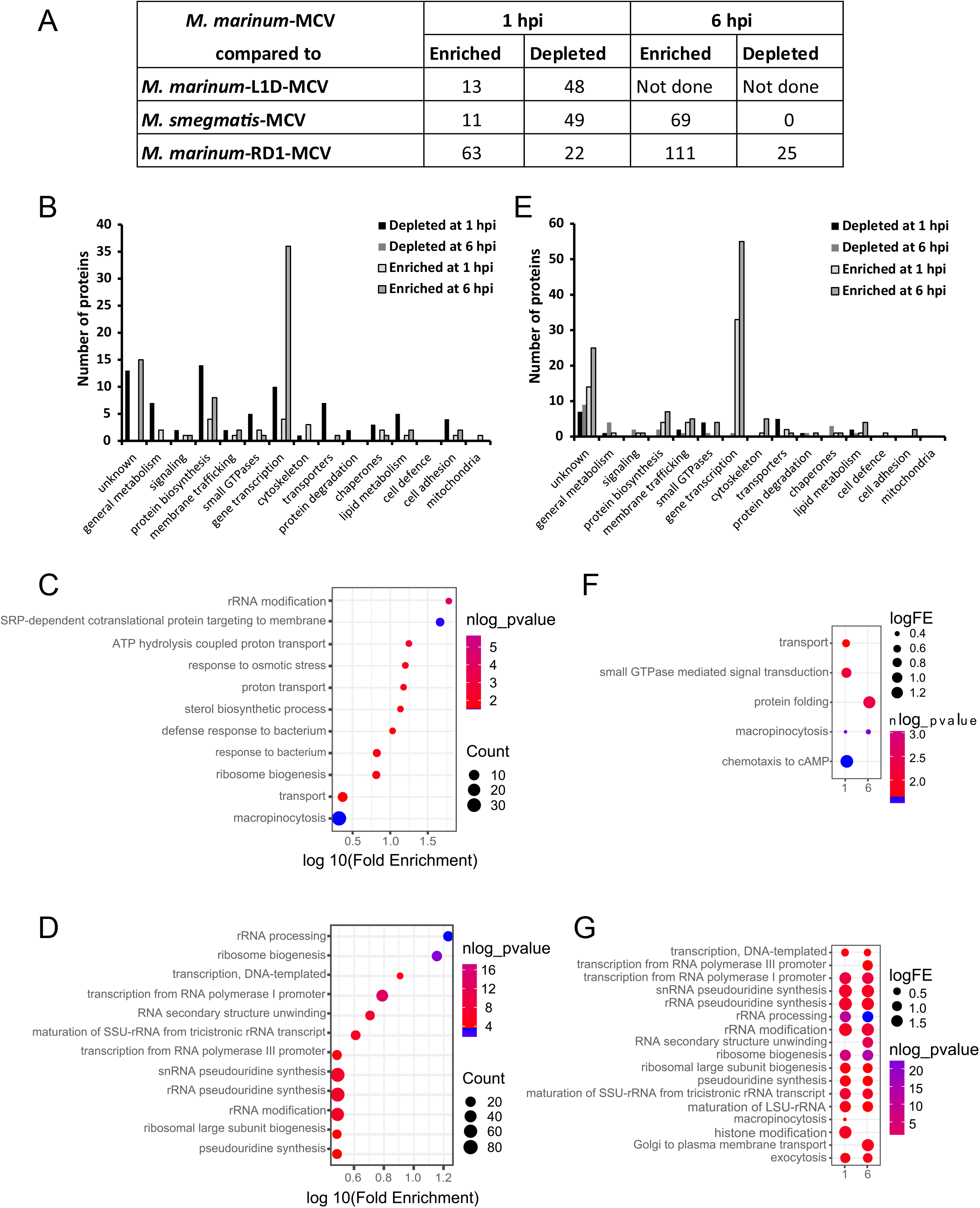
*M. marinum* MCV comparison with nonmanipulated Mycobacterium-containing compartments. MCVs containing *M. marinum*, *M. marinum*-L1D, *M. smegmatis* or *M. marinum*-RD1 were isolated at 1 and 6 hpi. All samples were labelled with TMT. *M. marinum*-MCVs were mixed with equal amounts of each other mycobacteria-MCVs. The obtained pools were analysed and quantified by LC-MS/MS. For each identified protein, the protein ratios of the corresponding comparison were calculated. **A**. The numbers of significantly differentially abundant proteins for each comparison are summarized in the table. *M. marinum*-MCV proteins significantly differentially abundant from *M. marinum*-L1D-MCV and/or *M. smegmatis*-MCV **(B)** or from *M. marinum*-RD1-MCV **(E)** were functionally classified using the same classification terms as in Figure 4. *M. marinum*-MCV proteins significantly depleted at 1 hpi **(C)** or enriched at 6 hpi **(D)** compared to *M. marinum*-L1D-MCV and/or *M. smegmatis*-MCV, or significantly depleted **(F)** or accumulated **(G)** compared to *M. marinum*-RD1-MCV were functionally annotated using the DAVID functional annotation clustering tool. The default parameters of the program were used and the category Gene Ontology “biological process” was selected. Enriched GO-terms are presented according to their enrichment score and p-value. Size of the bubble represents the number of proteins in the cluster.

At 1 hpi already, numerous proteins were found differentially abundant indicating that as early as at 1 hpi, *M. marinum* has diverted its compartment from the normal phagosome maturation pathway. Furthermore, the 1 hpi comparisons results reflect the different fate of the mycobacteria strains used: *M. marinum*-L1D and *M. smegmatis* are not able to manipulate their containing-vacuole and are rapidly killed and exocytosed whereas *M. marinum*-RD1, despite lacking the RD1 locus necessary to manipulate phagosome maturation, can survive in its MCV up to 30 hours, indicating a partial manipulation of its containing-vacuole (Cardenal-Muñoz et al., 2017). Indeed, at 1 hpi, *M. marinum*-MCV is mainly depleted in proteins compared to *M. marinum*-L1D and *M. smegmatis*-MCV, whereas it is mainly enriched in proteins compared to *M. marinum*-RD1-MCV. For this reason, *M. marinum*-MCV will first be compared to *M. marinum*-L1D and *M. smegmatis*-MCVs. The comparison of *M. marinum*-MCV with *M. marinum*-RD1-MCV will be discussed later. For both comparisons, proteins accumulating or depleted from *M. marinum*-MCV were manually sorted (**Figure 6B, E)** and functionally annotated using the DAVID functional annotation clustering tool (**Figure 6C, D, F, G**) (Huang et al., 2009; Sherman et al., 2022).

Not surprisingly, GO terms “defence response to bacterium” and “response to bacterium” were significantly enriched among the proteins depleted from the *M. marinum*-MCV at 1 hpi, indicating the MCV has already diverted from the normal phagosomal maturation pathway to become a non-degradative compartment. The high representation of the classes “signalling”, “membrane trafficking”, “small GTPases” and “transporters” and the significant enrichment of the GO terms “transport”, “proton transport”, “ATP hydrolysis coupled proton transport” and “response to osmotic stress” among the proteins depleted at 1 hpi indicates that this diversion results from the arrest or at least the limited interaction of the *M. marinum*-MCV with other endocytic compartments, and notably lysosomes. Among the proteins less abundant on the *M. marinum*-MCV at 1 hpi were notably found proteins involved in intracellular bacterial killing like Phg1a and Kil1 (Benghezal et al., 2006; Le Coadic et al., 2013), vATPase subunits necessary for intraphagosomal acidification (Clarke et al., 2002), the lysosomal enzyme Cystein protease, proteins involved in membrane trafficking like annexin 7 and vps13c and several small GTPases involved in vesicular trafficking and fusion like Sar1A, Rab14, Rab2a, an Arf GTpase and the GTPase binding protein Yip1. Numerous proteins involved in gene transcription and mRNA translation (classes “protein biosynthesis” and “gene transcription and GO terms “rRNA modification” and “ribosome biogenesis”) were also found depleted in the *M. marinum*-MCV, indicating again its limited fusion with other intracellular compartments, and notably autophagosomes.

Interestingly, the protein class “lipid metabolism” (Dgat2, Erg24, Lip5, sterol 14-demethylase, SmtA) and the GO term “sterol biosynthetic process” were found highly depleted from the *M. marinum*-MCV at 1 hpi. These proteins might have relocalised to lipid droplets in order to favour production and storage of lipids. Indeed, lipid droplets rapidly cluster around the *M. marinum*-MCV after infection. Their content is then transferred into the replicative niche (Barisch et al., 2015).

Among the proteins enriched on the *M. marinum*-MCV at 1 hpi, the class “cytoskeleton” was found: LimE, PirA and WH2 domain-containing protein. Two small GTPases were also found enriched: RasB and Arf1. Interestingly, Arf1, after activation by ACAP-A is involved in actin cytoskeleton remodelling. The enrichment of both cytoskeleton proteins and Arf1 confirms that the *M. marinum*-MCV retains an actin coat to prevent delivery of the vATPase (Kolonko et al., 2014). The retromer subunit Vps5 was also enriched. The retromer, through its interaction with the WASH complex, might be involved in recycling processes both at early (surface proteins) and late (mannose-6-phosphate tagged lysosomal enzymes) macropinosome stages (Buckley et al., 2016). Its enrichment on *M. marinum*-MCV at 1 hpi could indicate an arrest of early recycling steps in order to keep early phagosomal marker and consequently, its accumulation on the MCV. It could also indicate an active retrieval of lysosomal enzymes.

At 6 hpi, on the *M. marinum*-MCV, we found only enriched proteins compared with *M. smegmatis*-MCV. A vast majority of these proteins were involved in gene transcription and mRNA translation as indicated by the high representation of the protein classes “protein biosynthesis” and “gene transcription” and the significantly enriched GO terms, all related to transcription and translation machineries. This confirms the non-degradative environment in the *M. marinum*-MCV, allowing the accumulation of proteins potentially targeted to the phagosome for degradation. Despite being slightly represented, two other proteins classes were found enriched on the *M. marinum*-MCV at 6 hpi: “membrane trafficking” and “lipid metabolism”. The two proteins involved in lipid metabolism were PLC and oxysterol binding protein 7 (OSBP7). At this stage, it is known that lipid droplet content can be observed in the *M. marinum*-MCV (Barisch et al., 2015). Host lipids are then used by the mycobacteria. Oxysterol binding protein 7 could be important for the transfer of lipids from the lipid droplets to the MCV. PLC would allow *M. marinum* to use host phospholipids as previously proposed (Barisch and Soldati 2017). Interestingly, the two enriched proteins involved in membrane trafficking were subunits of the exocyst tethering complex (Exoc2 and Exoc1).

The RD1 locus is known to be necessary for the establishment of a successful infection. It encodes the ESX-1 secretion system, essential to induce damages to the MCV membrane (Cardenal-Muñoz et al., 2017; López-Jiménez et al., 2018). At later times of infection, after intra-MCV replication, those damages allow mycobacteria to escape the MCV, access nutrient into the cytosol and disseminate to neighbouring cells. However, early escape from the MCV, before replication, would be deleterious for the infection. In the cytosol, the bacterium would be captured by autophagy and targeted to lysosome for degradation (Cardenal-Muñoz et al., 2017). To prevent this early escape, the ESCRT and the autophagy machineries are both recruited to the MCV in the form of patches to seal the injured membrane (López-Jiménez et al., 2018). As a consequence, it has been shown that the *M. marinum*-RD1 mutant survives in its MCV but is not able to arrest phagosome maturation (Tan et al., 2006), to recruit lipid droplets around its MCV (Barisch et al., 2015), cannot induce damages nor escape from its MCV (Cardenal-Muñoz et al., 2017; López-Jiménez et al., 2018), cannot replicate and cannot form ejectosomes to disseminate to neighbouring cells (Hagedorn et al., 2009).

When compared to *M. marinum*-RD1-MCV, only few proteins were found depleted from the *M. marinum*-MCV, both at 1 and 6 hpi. Both manual protein sorting and GO annotation indicate that at 1 hpi, those proteins are small GTPases and transporters. Several of those transporters (RghA, Nhe1) are involved in osmoregulation. Their enrichment on the *M. marinum*-RD1-MCV could reflect the potentially acidic and toxic ionic environment in its lumen. The small GTPases accumulated on *M. marinum*-RD1-MCV (RasG, RacE, RasS) have been shown to be involved in actin cytoskeleton remodelling in pinocytosis, cytokinesis and cell motility. As numerous other small GTPases, a function in endosome trafficking cannot be excluded. Their enrichment on the *M. marinum-RD1*-MCV could indicate that the compartment follows the normal phagosomal maturation pathway.

Numerous proteins were found accumulated at the *M. marinum*-MCV compared to the *M. marinum*-RD1-MCV. Both at 1 and 6 hpi, the protein classes “gene transcription” and “protein biosynthesis” were strongly represented. Consistent with that, almost all the significantly enriched GO-terms were related to transcription machinery. This again indicates the non-degradative environment of the *M. marinum*-MCV lumen and consequently the accumulation of cytoplasmic material delivered by autophagosomes. On the contrary, the depletion of this material from the *M. marinum*-RD1-MCV confirms that the RD1 locus is necessary to induce damages to the MCV and the following recruitment of the autophagic machinery, essential to the repair of those membrane damages (Cardenal-Muñoz et al., 2017; López-Jiménez et al., 2018). The protein class “lipid metabolism” was also enriched in the *M. marinum*-MCV at 6 hpi. OSBP7 is notably found in this class. As previously mentioned, OSBP7 could be involved in the transfer of lipid droplet content to the MCV. Recently, it has been shown that oxysterol binding proteins are also involved in lysosomal membrane damage repair by forming contact sites between the damaged compartment and the ER. They provide lipids for the repair by exchanging PI4P in excess in the damaged membrane with cholesterol from the ER (Radulovic et al., 2022; Tan and Finkel, 2022). OSBP7 could play here a similar function and participate to the MCV membrane damage repair. Consistent with that, participation of OBPB8 to *M. marinum*-MCV membrane repair has recently been demonstrated (Anand et al., 2023). The other enriched protein class was “membrane trafficking” and the other significantly enriched GO terms were “exocytosis” and “Golgi to plasma membrane transport”. Interestingly, proteins found in those classes and GO terms are subunits of the exocyst complex. Four subunits were enriched at 1 hpi (Exoc1, Exoc2, Exoc3 and Exoc7) and five at 6 hpi (Exoc1, Exoc2, Exoc3, Exoc4 and Exoc5).

In conclusion, comparison of *M. marinum*-MCV to non or less manipulated MCVs confirmed the previously observed 2 phases in the establishment of a replicative niche. During the first phase, the *M. marinum*-MCV has very limited interaction with other endosomal compartments, and especially with late endosomes and lysosomes. This is notably permitted by the retained actin coat (Kolonko et al., 2014). Furthermore, according to the early *M. marinum*-MCV maturation analysis, fission seems to be blocked as well, preventing the removal of early phagosomal markers and further maturation. However, recycling of specific proteins like lysosomal enzymes might still occur, as vps5, a subunit of the retromer complex accumulates on the *M. marinum*-MCV (Buckley et al., 2016). As a result, *M. marinum*-MCV do not acidify, do not acquire lysosomal enzyme nor proteins involved in bacterial killing, and its lumen remains no degradative. During the first phase, the *M. marinum*-MCV is also depleted in proteins involved in lipids transport and metabolism. Those proteins might be relocalised to lipid droplets to favour lipid storage in those compartments. During the second phase, *M. marinum*-MCV starts interacting again with non-degradative intracellular compartments, and notably autophagosomes (Cardenal-Muñoz et al., 2017). As a result, non-digested cytoplasmic material accumulates in its non-degradative lumen. The autophagic machinery recruited during this second phase also allow to repair the MCV membrane damages induced by *M. marinum* and more precisely by the proteins encoded by its RD1 locus. Furthermore, proteins involved in lipid metabolism accumulate on the *M. marinum*-MCV during the second phase. This allows the mycobacteria to access and use host lipids (Barisch et al., 2015; Barisch and Soldati, 2017). Lipids transporters like OSBP7 would also participate to the MCV membrane damage repair by providing lipids.

### SnxA, a new early-MCV marker

Only one protein was found constantly accumulating on the *M. marinum*-MCV during the manipulation phase: a phox domain-containing protein. We decided to focus on this protein as it was recently identified as a very specific PI(3,5)P_2_ probe and named SnxA (Vines et al., 2022). We will hereafter also refer to this phox domain-containing protein as SnxA.

*D. discoideum* cells expressing SnxA-GFP were infected with *M. marinum* and observed by live-microscopy. Confirming the quantitative proteomic results, the percentage of SnxA positive MCV was increasing during the manipulation phase from 38.0 % at 1 hpi to 55.6 % at 6 hpi (**Figure 7A, B)**. However, this SnxA accumulation is transient and restricted to the manipulation phase. Indeed, as early as 12 hpi, this percentage dropped to 26.8 %, below the percentage measured at the earliest time of infection (**Figure 7C, D**). 12 hpi corresponds to the transition from the manipulation phase to the proliferation phase. During this second phase, the MCV becomes spacious and *M. marinum* starts proliferating. Consistent with that, spacious SnxA-positive MCV were rarely observed while SnxA-positive MCV, with the membrane tightly apposed to the mycobacteria were mainly observed (**Figure 7A, C**). As SnxA is a PI(3,5)P_2_ probe, this result indicates that the *M. marinum*-MCV membrane contains PI(3,5)P_2_ during the manipulation phase. As PI(3,5)P_2_ is mainly found in the membrane of late phagosomes and lysosomes, its loss from the MCV membrane might indicate the final transition from a phagosomal compartment to a replicative niche. Indeed, this PI(3,5)P_2_-negative MCV no longer interacts with late acidic endosomes, nor lysosomes. *M. marinum* resides in a non-stressful environment where it can finally start to replicate.

**Figure 7.**
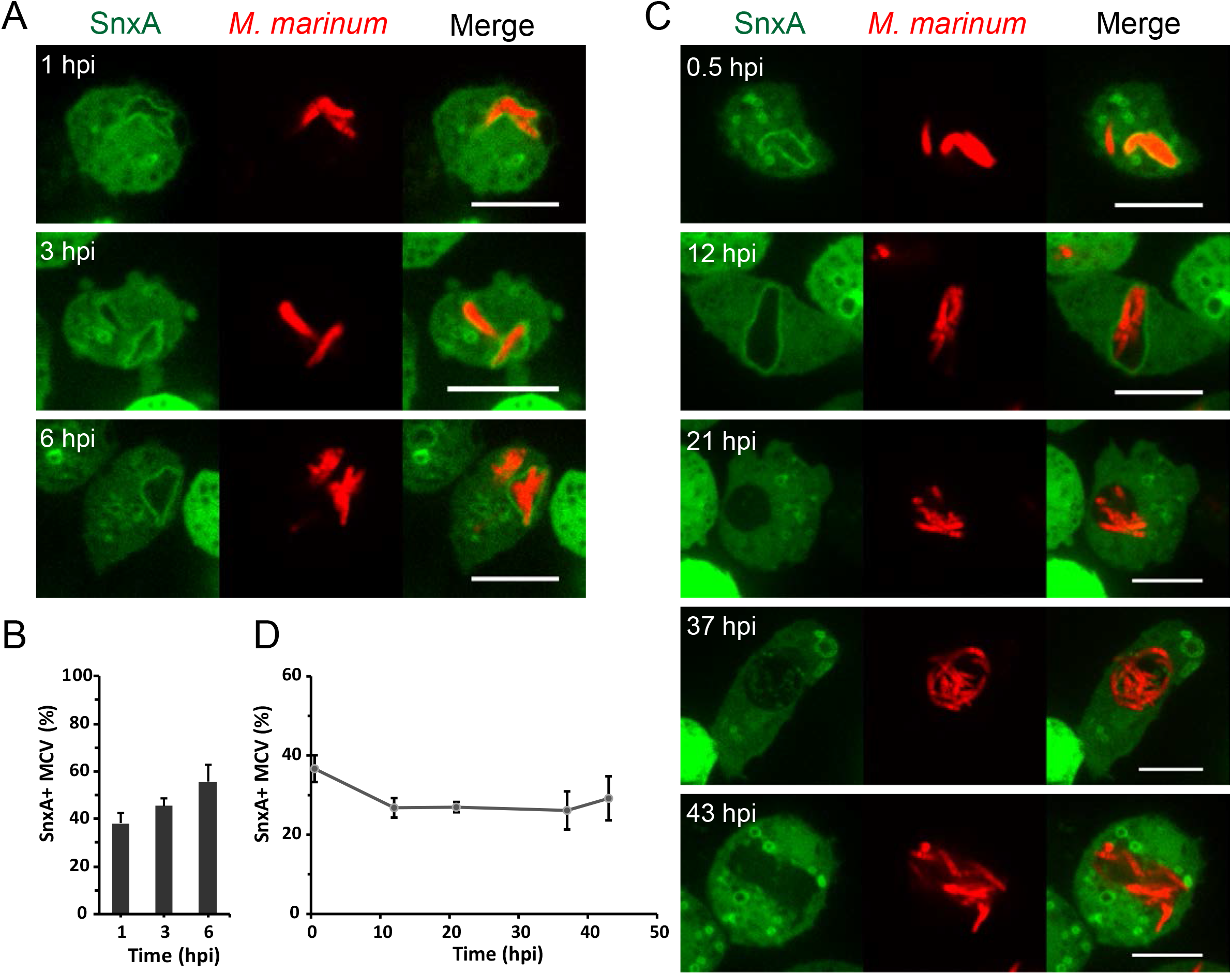
SnxA is an early M. marinum-MCV marker. SnxA-GFP expressing cells were infected with M. marinum-RFP. At early infection times (1, 3 and 6 hpi), cells were observed by live-microscopy. Scale bars, 10 μm. (**A**). The number of SnxA-positive MCVs was counted. Graphs represent mean and SEM of 3 independent experiments (**B**). SnxA-GFP expressing cells infected with M. marinum-RFP were observed by live microscopy at 0.5, 12, 21, 37 and 43 hpi. Scale bars, 10 μm (**C**). The number of SnxA-positive MCVs was counted. Graphs represent mean and SEM of 3 independent experiments (**D**).

SnxA is thus a new MCV marker that accumulates in the *M. marinum*-MCV specifically during the manipulation phase. Its removal from the MCV can help visualizing the transition from the manipulation to the replicative phase.

### A role for the exocyst in MCV maturation

The exocyst complex is a tethering complex involved in exocytic processes, and notably vesicle trafficking. It is composed of 8 subunits which are separated in two subcomplexes. In *D. discoideum*, the first subcomplex would be composed of Sec3 and maybe Exo84, whereas the second subcomplex would be composed of Sec15, Sec6, Sec8, Sec10, Exo70 and potentially Sec5 (Essid et al., 2012). Interestingly, the exocyst subunits were all identified in the early *M. marinum*-MCV. The subunit Sec8 seems to be transiently depleted at 3 hpi during the *M. marinum*-MCV maturation. Despite being not significant, the ratios measured for the other subunits also tend to confirm a transient depletion of the exocyst from *M. marinum*-MCV at 3 hpi, followed by an accumulation to return at 6 hpi to level similar as at 1 hpi. Furthermore, six of the exocyst complex subunits (Sec3, Sec5, Sec6, Sec8, Sec 10 and Exo70) were also differentially abundant in the comparison between the *M. marinum*-MCV and the “non-manipulated”-MCVs (**Figure 8A**). They were enriched in *M. marinum*-MCV compared to *M. marinum*-RD1-MCV both at 1 and 6 hpi, and compared to *M. smegmatis*-MCV at 6 hpi. Again, for some subunits, the measured ratios were not considered significant, but were consistent with an enrichment of the exocyst in the *M. marinum-MCV*. We decided to focus on the subunit Sec8 since it was significantly differentially abundant both in the early *M. marinum*-MCV maturation analysis and in the comparison of *M. marinum*-MCV to “non manipulated”-MCVs.

**Figure 8.**
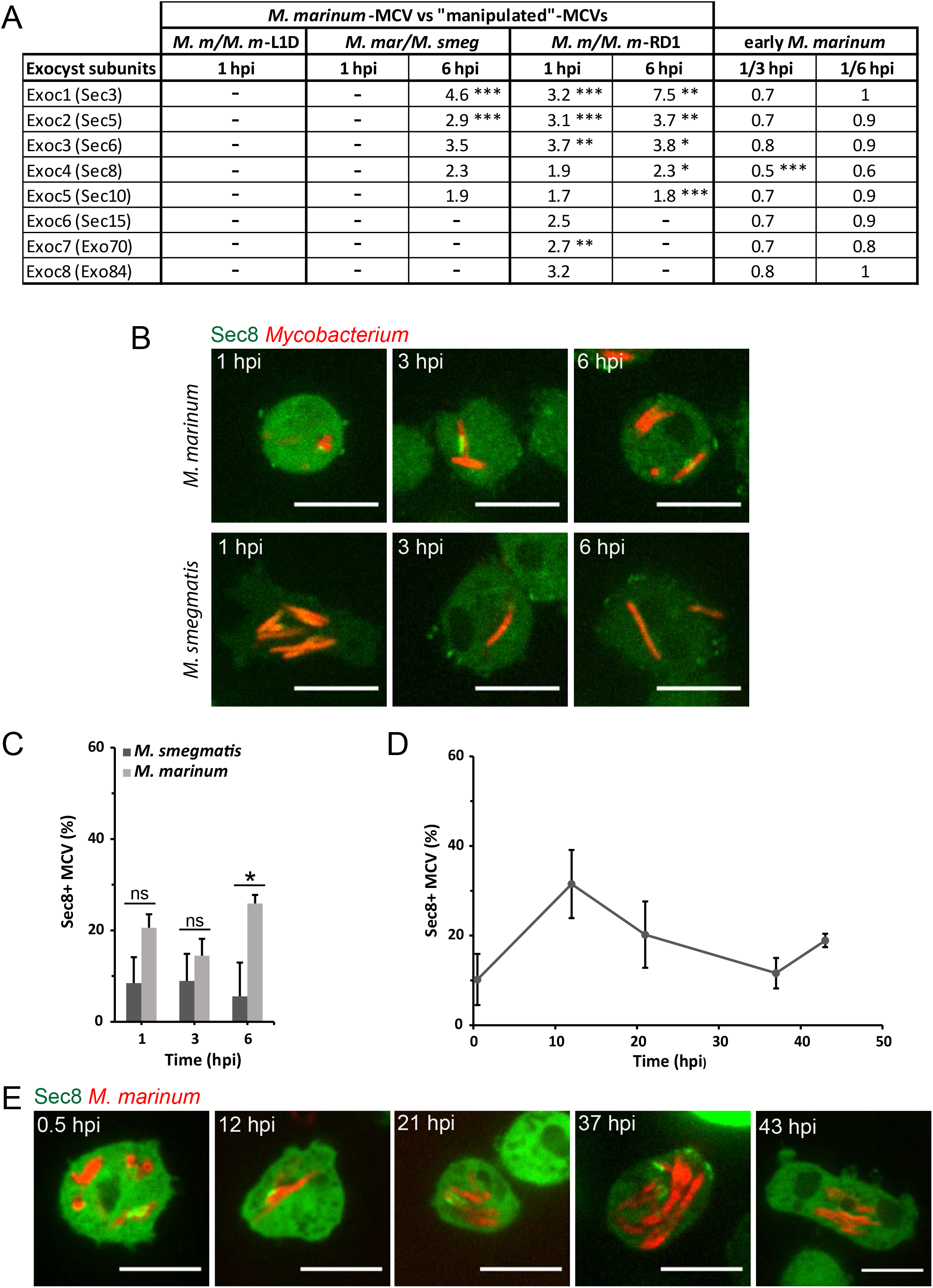
The exocyst in MCV maturation. Ratios calculated for each exocyst subunits in the different proteomic comparison performed: M. marinum/”non manipulated” MCVs and M. marinum-MCVs at 1 hpi/M marinum-MCVs 3 hpi or 6 hpi (ns = not significant) (**A**). Sec8-GFP expressing cells were infected with M. marinum-RFP or M. smegmatis-DsRed. At early infection times (1, 3 and 6 hpi), cells were observed by live-microscopy. Scale bars, 10 μm. (**B**). The number of Sec8-positive MCVs was counted. Graphs represent mean and SEM of 3 independent experiments. Two-ways ANOVA followed by a Tukey’s test was performed (**C**). Sec8-GFP expressing cells infected with M. marinum-RFP were observed by live microscopy at 0.5, 12, 21, 37 and 43 hpi. Scale bars, 10 μm (**E**). The number of SnxA-positive MCVs was counted. Graphs represent mean and SEM of 3 independent experiments (**D**).

*D. discoideum* cells expressing Sec8-GFP were infected with *M. marinum* or *M. smegmatis* and observed by live-microscopy. Sec8 recruitment to MCVs consisted in patches or dots along mycobacteria. MCVs entirely Sec8 positive were not observed (**Figure 8B**). Confirming the quantitative proteomic results, the percentage of Sec8 positive *M. marinum*-MCV was slightly decreased at 3 hpi. Furthermore, this percentage was higher for cells infected with *M. marinum* than for cells infected with *M. smegmatis*. At 6 hpi, this difference was significant (**Figure 8C**). On the entire infection cycle of *M. marinum*, a pic of 31.5 % of Sec8 positive *M. marinum*-MCVs was observed at 12 hpi. As suggested earlier, this confirms that after a transient depletion, Sec8 accumulates in *M. marinum*-MCV from 3hpi. After 12 hpi, the percentage of Sec8 positive *M. marinum*-MCV progressively decreased (**Figure 8D, E**).

The exocyst complex has been shown to be involved in the establishment of a successful infection of other intraphagosomal pathogens. *Legionella pneumophila* remodels its phagosome-like containing compartment into an ER-like compartment by promoting its fusion with ER-derived vesicle. Sec5 and Sec15 participate to the assembly of the SNARE complex, allowing the recruitment of those ER-derived vesicles to the *Legionella* containing vacuole (Arasaki et al., 2018). *Shigella* recruits infection-associated macropinosomes (IAM) around its containing-vacuole. Those IAM promote the bacteria escape into the cytosol. The exocyst is involved in this IAM clustering step (Chang et al., 2020). As *Legionella* and *Shigella*, *M. marinum* might hijack the exocyst for the recruitment of vesicles or protein complex to its MCV.

The enrichment of the exocyst in *M. marinum*-MCV compared to *M. marinum*ΔRD1-MCV both at 1 and 6 hpi suggests that the exocyst recruitment might depend on the RD1 locus. As previously mentioned, this locus is essential to induce damages to the MCV membrane and allow mycobacteria to escape into the cytosol and disseminate to neighbouring cells at later times of infection. However, to prevent an early escape which would be deleterious for the infection, host protein complexes of the ESCRT and the autophagy machineries are both recruited to the MCV in the form of patches to seal the injured membrane(Cardenal-Muñoz et al., 2017; López-Jiménez et al., 2018). Interestingly, Bodeman et al. have shown that the exocyst complex interacts with the autophagy machinery in human epithelial cells, and is involved in the autophagosome membrane nucleation and elongation (Bodemann et al., 2011). The exocyst might have a similar function during *M. marinum* infection in *D. discoideum*. Consistent with this hypothesis, the recruitment of the exocyst subunit Sec8 to the *M. marinum*-MCV is very similar to the dotty and patchy Atg8 recruitment previously observed in *D. discoideum* infected with *M. marinum* (**Figure 8A, C**) (Cardenal-Muñoz et al., 2017; López-Jiménez et al., 2018). However, the different subunits of the exocyst do not contribute similarly to the autophagosome biogenesis. Indeed, upon autophagy induction signal, the inactive autophagosome biogenesis machinery disassemble from the Sec5 subunit to reassemble catalytically active on Exo84 (Bodemann et al., 2011). The individual function and contribution of the different exocyst subunits to autophagy and to the *M. marinum* infection in *D. discoideum* would require further investigations.

In this study, we established a new procedure to isolate pure fractions of MCVs. This allowed us to isolate and characterize the proteome of the early *M. marinum*-MCV during the manipulation phase of the infection process. Then, using differential analysis of *M. marinum*-MCVs isolated at 1, 3 and 6 hpi, 2 phases could be distinguished during the manipulation of the containing-compartment to tailor it into a replicative niche. A second proteomic comparison of *M. marinum*-MCVs with non or less-manipulated MCVs isolated at 1 and 6 hpi confirmed that the manipulation phase of the infection process was divided into two distinct phases, and helped to further dissect the processes occurring during those phases.

The first phase consists in arresting the maturation of the compartment into a degradative phagosome. During this phase, the *M. marinum*-MCV retains an actin coat which prevents it from interacting with other compartments of the endo-lysosomal pathway (Kolonko et al., 2014). Furthermore, fission is partially blocked. This allows the *M. marinum*-MCV to retain early phagosomal markers, and prevent the compartment to process to the next step of phagosome maturation. However, the presence of proteins like the vps5 subunit of the retromer complex indicates that recycling of some specific protein like lysosomal enzymes might still occur (Buckley et al., 2016). Proteins involved in lipid transport and metabolism are relocalised to lipid droplet to store lipids which will be used latter by *M. marinum*.

The second phase consists in maintaining the membrane integrity of the replicative niche. During this phase, *M. marinum*-MCV resumes interactions with non-degradative intracellular compartments and notably autophagosomes (Cardenal-Muñoz et al., 2017). As a result, undigested cytoplasmic materials accumulate in its non-degradative lumen. Furthermore, through proteins encoded by its RD1 locus, *M. marinum* induces damages to its containing compartment. The autophagic machinery recruited to the *M. marinum*-MCV during this phase actively seals those damages (Cardenal-Muñoz et al., 2017). Finally, proteins involved in lipid metabolism accumulate in the *M. marinum*-MCV to allow the mycobacterium to access and use host lipids (Barisch et al., 2015; Barisch and Soldati, 2017). Some lipid transporters like OSBP7 might also have a role in the *M. marinum*-MCV membrane repair (Anand et al., 2023; Radulovic et al., 2022; Tan and Finkel, 2022).

Our differential proteomic analyses also allowed us to identify a new early *M. marinum*-MCV marker: SnxA. This PI(3,5)P_2_ probe transiently accumulates to the *M. marinum*-MCV during the manipulation phase (Vines et al., 2022). As PI(3,5)P_2_ is mainly found in the membrane of late phagosomes and lysosomes, its loss from the *M. marinum*-MCV might reflect the final diversion step of the compartment from a phagosome into a replicative niche. SnxA could then be used as a new tool to visualize this transition.

Finally, our results indicate a potential new role for the exocyst complex in the establishment of a successful mycobacterial infection. It is transiently recruited to the *M. marinum*-MCV during the first twelve hours of infection. According to our proteomic results, its recruitment seems to be RD1-dependent. Furthermore, it is involved in autophagosome membrane nucleation and elongation (Bodemann et al., 2011). It is then tempting to imagine a role for the exocyst complex in the MCV membrane damage repair through its interaction with the autophagy machinery. This potential new function would require further investigations.

## MATERIAL AND METHODS

### *D. discoideum* cell culture

Wild-type *D. discoideum* cells (AX2) were cultivated at 22°C in HL5c medium supplemented with 100 U/mL penicillin and 100 μg/mL streptomycin (Invitrogen). The *lmpB* ko cells were kindly provided by Dr M. Schleicher (Janssen et al., 2001; Karakesisoglou et al., 1999). AX2 cells expressing SnxA-GFP were described in (Vines et al., 2022). AX2 cells expressing Sec8-GFP were described in (Essid et al., 2012).

### Mycobacteria culture

Mycobacteria were cultivated at 32°C under shaking conditions (150 rpm) in Middlebrook 7H9 medium (Difco), 0.2% glycerol and 0.05% Tween 80 supplemented with 10 % OADC (Beckton Dickinson). They were cultivated in presence of 5 mm glass beads in order to avoid bacteria clumping. *M. marinum*-GFP, *M. marinum* RD1-GFP and *M. smegmatis*-GFP and *M. smegmatis*-DsRed were cultivated in presence of 50 μg/mL kanamycin, *M. marinum* L1D-GFP in presence of 50 μg/mL apramycin and *M. marinum*-LuxABCDE and mCherry-expressing *M. marinum* in presence of 50 μg/mL and 100 μg/μL hygromycin, respectively.

### Preparation of bacteria+beads complexes (BBCs) inoculum

Latex beads and 5.10^9^ mycobacteria were washed 3 times separately in 0.1 M borate for 10 min at 12 000 rpm, RT. The amount of latex beads to prepare was adapted according to the latex bead size: beads:bacteria ratio of 500:1 for 0.2 μm beads, 90:1 for 0.5 μm beads and 15:1 for 0.8 μm beads. After resuspension in 0.1 M borate, latex beads were sonicated for 5 min in a sonicator bath. The latex beads and the mycobacteria were then mixed and incubated on a rotating wheel for 2 h. The mycobacteria+beads mix was subsequently washed once in HL5c medium for 10 min at 12 000 rpm, and resuspended in HL5c. In order to remove unattached beads, the mix was deposited on top of a one-step sucrose gradient (20% sucrose and 60% sucrose) and ultracentrifuged for 30 min at 55 000 rpm in a TLS55 rotor. The pure inoculum of mycobacteria+beads complexes was recovered at the 20-60% sucrose interphase. It was then washed once in HL5c medium for 10 min at 12 000 rpm and finally resuspended in HL5c medium without pen/strep.

### Phagocytosis assay

Phagocytosis of *M. marinum*-GFP or *M. marinum*-GFP-BBCs by AX2 and *LmpB ko* cells was monitored by FACS as previously described (Sattler et al., 2013). Briefly, cells from confluent plates were resuspended in 5 mL of fresh HL5c medium at a density of 2.10^6^ cells/mL and agitated for 2 h at 150 rpm, 22°C, prior to the experiment. 2.10^8^ *M. marinum*-GFP were prepared per cell line (MOI = 100). Bacteria were washed twice in HL5c medium and resuspended in 1mL of HL5c. In order to declump aggregates, mycobacteria were passaged through a 25-gauge needle for 8-10 times. 0.2 μm *M. marinum*-GFP-BBCs were prepared as described before. At T0, cells were agitated at 120 rpm and the mycobacteria or BBCs were added to the cells. At each time point, an aliquot was taken. The phagocytosis was stopped by adding one volume of ice-cold Sorensen-Sorbitol-Azide (15 mM KH_2_PO_4_, 2 mM Na_2_HPO_4_, 120 mM Sorbitol, 5 mM azide). After centrifugation 10 min at 1200 rpm at 4°C, cells were resuspended in Sorensen-Sorbitol (15 mM KH_2_PO_4_, 2 mM Na_2_HPO_4_, 120 mM sorbitol) and the fluorescence was measured by FACS. FACS measurement was performed on a FACScalibur (Beckton Dickinson) and the data analysed with FlowJo (TreeStar, USA).

### Infection assay

Infections were performed as previously described (Arafah et al., 2013; Hagedorn and Soldati, 2007; Sattler et al., 2013). *D. discoideum* cells were infected with 5.10^8^ mycobacteria or 500 μL of prepared BBCs inoculum. Infections with GFP-expressing mycobacteria were monitored by FACS (Sattler et al., 2013). At times of interest, infection aliquots were mixed V:V with Sorensen-azide (15 mM KH2PO4, 2 mM Na2HPO4, 5 mM azide) and pelleted for 4 min at 12 000 rpm, RT. The pellet was resuspended in Sorensen-Sorbitol (15 mM KH2PO4, 2 mM Na2HPO4, 120 mM sorbitol). Prior to FACS measurement, 4.5 μm fluorescent beads (100 beads/μL, YG-beads, Polyscience) were added to the sample as an internal particle concentration standard. FACS measurement was performed on a FACScalibur (Beckton Dickinson) and the data analysed with FlowJo (TreeStar, USA). Infections with bioluminescent bacteria were monitored with a plate reader (Synergy Mx, Biotek) (Arafah et al., 2013). At times of interest, 150 μL aliquots of infected cells were deposited in a white 96-well plate (F96 MicroWell™ Plates, non-treated form Nunc) and the luminescent signal was measured with the help of the plate reader.

### Immunofluorescence

*D. discoideum* infected cells and BBCs were fixed with PFA/picric acid or by shock-freezing fixation in −85°C methanol as described previously (Hagedorn et al., 2006). The following antibodies were used: vacuolin (221-1-1, (Dr. M. Maniak (Jenne et al., 1998)), VatM (N2, Dr. R. Allen (Fok et al., 1993)), mitochondrial porin (Dr. G. Gerisch), p80 (purchased from the Geneva Antibody Facility), cocktail *M. leprae* MLSA, MLMA and MLcwA, calreticulin (mAb, 251-67-1) (gift of Dr. G. Gerisch, MPI for Biochemistry, Martinsried), PDI. Goat anti-mouse or anti-rabbit IgG coupled to AlexaFluor594 (Invitrogen) were used as secondary antibodies.

### EM

Cells were fixed and stained as previously described, (Barisch et al., 2015). Cells were fixed for 1 h in 2% glutaraldehyde and then, stained for 30 min in a 2% osmium/0.1 M imidazole solution. After 2 washes in PBS, cells were then sent to the EM platform of the Faculty of Medicine, University of Geneva, where the samples were fixed, embedded in Epon resin and processed for EM. Grids were observed with a Tecnai 20 electron microscope (FEI).

### Live fluorescence microscopy

Infected *D. discoideum* cells were plated in 35 μ-dish (Ibidi, Gräfelfing, Germany). Images were acquired with a Spinning Disc Confocal Microscope (Intelligent Imaging Innovations, Göttingen, Germany) mounted on an inverted microscope (Leica DMIRE2; Leica, Wetzlar, Germany) with a glycerin immersion 63x objective. Image analysis was performed using ImageJ.

### Isolation of MCV

At time of interest (1, 3 or 6 hpi), the infection was stopped by pelleting the infected cells at 4°C for 8 min at 2000 rpm. Cells were resuspended in cold HESES-azide buffer (20 mM HEPES-KOH, 0.25 M sucrose, 5 mM azide, pH 7.2) containing a proteaseinhibitor cocktail (Complete EDTA-free, Roche) and homogenized by passing them 8 times through a ball homogenizer (HGM, Germany) with a 10 μm clearance. The cell homogenate was then incubated with 10 mM Mg-ATP on a rotating wheel for 15 min at 4°C and loaded onto sucrose step gradients (layers of 10%, 30%, 40% and 60% sucrose). After overnight ultracentrifugation of the gradients at 28 000 rpm in a SW40 rotor at 4°C, MCVs were collected at the 10-30% sucrose interphase and diluted in membrane buffer (20 m*M* HEPES-KOH, pH 7.2, 20 m*M* KCl, 2.5 m*M* MgCl2, 1 m*M* dithiothreitol (DTT), 20 m*M* NaCl). MCVs were then pelleted by ultracentrifugation for 1h30 at 35 000 rpm at 4°C in a SW40 rotor. Dry pellets of MCV were finally snap-frozen and stored at −80°C until further analysis.

### Samples reduction, alkylation, digestion and TMT labelling

The reduction, alkylation, digestion and TMT labelling were mainly performed as described by (Dayon et al., 2008). Briefly, the MCV pellets were resuspended in 50 μL of TEAB (Triethylammonium hydrogen carbonate buffer) 0.1 M pH 8.5, 6 M urea.

After addition of 50 mM of TCEP (tris-(2-carboxyethyl) phosphine hydrochloride), reduction was performed during 1 h at 37°C. The samples were then centrifuged several times for 10 min, at 12,000 rpm until the collected supernatants were completely free of latex beads. Protein concentration of each sample was measured with the help of a nanodrop. 100 μg of proteins of each sample were alkylated at RT in the dark for 30 min. after addition of 1 μL of IAA (Iodoacetamide) 400 mM. The volume of the samples was then adjusted to 100 μL with TEAB 0.1 M pH 8.5. After addition of 2 μg of trypsin, samples were digested overnight at 37°C. Each sample was then labelled with one TMT reagent according to manufacturer’s instructions. Finally, 50 μL of each labelled sample were pooled and evaporated under speed-vacuum.

### OGE

Off-gel electrophoresis was performed according to manufacturer’s instructions (Agilent). After desalting, the mix containing the pooled labelled samples was reconstituted in OFFGEL solution. A 12 or 24-wells frame was set-up on an Immobiline DryStrip pH 3-10, 24 cm and isoelectric focusing was performed with those settings: 8,000 V, 50 μA, 200 mW until 20 kVh was reached. The fractions were then recovered and desalted using C18 MicroSpin columns.

### Mass spectrometry

ESI LTQ-OT MS was performed on an LTQ Orbitrap Velos from Thermo Electron (San Jose, CA, USA) equipped with a NanoAcquity system from Waters. Peptides were trapped on a home-made 5 μm 200 Å Magic C18 AQ (Michrom) 0.1 × 20 mm pre-column and separated on a home-made 5 μm 100 Å Magic C18 AQ (Michrom) 0.75 × 150 mm column with a gravity-pulled emitter. The analytical separation was run for 65 min using a gradient of H2O/FA 99.9%/0.1% (solvent A) and CH3CN/FA 99.9%/0.1% (solvent B). The gradient ran as follows: 0–1 min 95% A and 5% B, then to 65% A and 61 35% B at 55 min, and 20% A and 80% B at 65 min at a flow rate of 220 nL/min. For MS survey scans, the OT resolution was set to 60000 and the ion population was set to 5 × 105 with an m/z window from 400 to 2000. A maximum of 3 precursors was selected for both collision-induced dissociation (CID) in the LTQ and high-energy C-trap dissociation (HCD) with analysis in the OT. For MS/MS in the LTQ, the ion population was set to 7000 (isolation width of 2 m/z) while for MS/MS detection in the OT, it was set to 2 × 10E5 (isolation width of 2.5 m/z), with resolution of 7500, first mass at m/z = 100, and maximum injection time of 750 ms. The normalized collision energies were set to 35% for CID and 60% for HCD.

### Protein identification

Protein identification was performed with the help of the Easyprot platform. The Easyprot platform proceeds as follows: peak lists are generated from raw data using (ReadW). After peaklist generation, the CID and HCD spectra are merged for simultaneous identification and quantification (Dayon et al., 2010) and (http://www.expasy.org/tools/HCD_CID_merger.html). The peaklist files were searched against the uniprot_sprot database (2011_02 of 08-Feb-2011). *Dictyostelium discoideum*, *Mycobacterium marinum* and *Mycobacterium smegmatis* taxonomies were specified for database searching. The parent ion tolerance was set to 10 ppm. TMT-sixplex amino terminus, TMT-sixplex lysine (229.1629 Da) and carbamidomethylation of cysteines were set as fixed modifications. Oxidized methionine was set as variable amino acid modification. Trypsin was selected as the enzyme, with one potential missed cleavage, and the normal cleavage mode was used. Protein and peptide scores were then set up to maintain the false positive peptide ratio below 1%. For identification, only proteins matching two different peptide sequences were kept.

### Protein quantification

Isobaric quantification was performed using the IsoQuant module of Easyprot’s protein export as described previously. Briefly, a false discovery rate of 1% and a minimum of 2 peptides per protein were selected. TMT sixplex was selected as reporter and a mass tolerance of 0.05 m/z was used. The different protein ratios were calculated. A global normalisation of “median log peptide ratio=0” and a confidence threshold of 95% were selected. EasyProt’s Mascat statistical method, inspired by Mascot’s quantification module (http://www.matrixscience.com/help/quant_statistics_help.html), and Libra, inspired by the Trans-Proteomic Pipeline’s isobaric quantification module (http://tools.proteomecenter.org/wiki/index.php?title=Software:Libra), were chosen, along the generation of the list of proteins featuring a ratio fold over 1.5. To determine the proteins differentially abundant, a cut-off threshold was calculated for each sample comparison as described previously (Tiberti et al., 2010). The results of those calculations are presented in **Table S5**.

---

**Figure S1.**
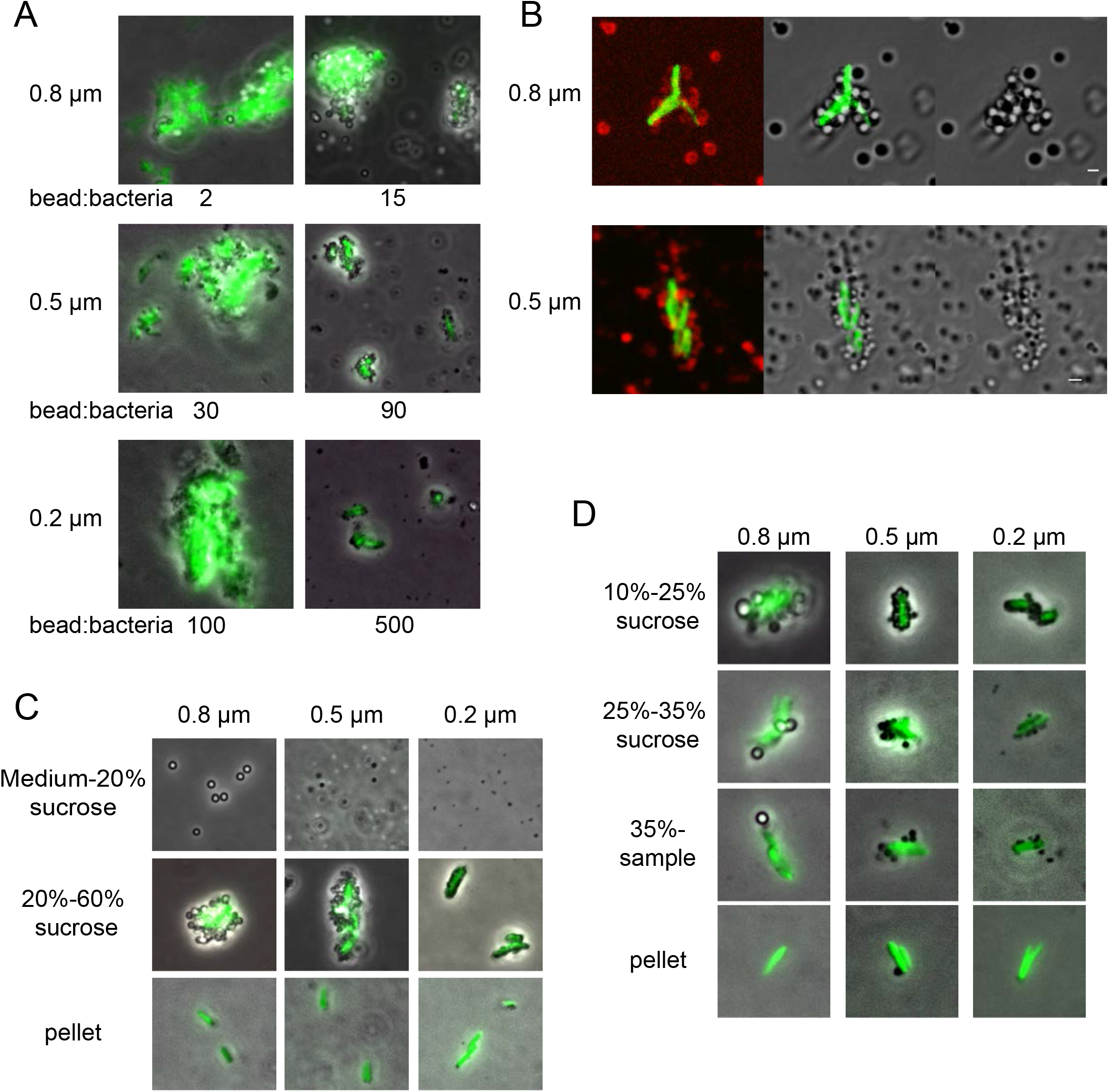
Latex beads adsorb onto Mycobacteria and can float them up on a sucrose gradient. **A**. *M. marinum*-GFP-BBCs were prepared with latex beads of different sizes and various beads:bacteria ratios. The resulting BBCs were visualized by microscopy. **B**. *M. marinum*-GFP-BBCs prepared with different latex bead sizes. TRITC-coupled antibodies were adsorbed onto latex beads during BBCs preparation. Scale bar, 1 μm. **C**. *M. marinum*-GFP-BBCs were prepared with 0.8 μm or0.5 μm latex beads at ratios 15 and 90 respectively. The prepared BBCs+free beads mixture was loaded onto a sucrose gradient. Fractions recovered at the different interphases were visualized by microscopy. **D**. *M. marinum*-GFP-BBCs prepared with 0.8 μm, 0.5 μm or 0.2 μm latex beads at ratios 15, 90 and 500 respectively and separated from free beads by ultracentrifugation on a one step sucrose gradient were loaded on the sucrose gradient used to isolate latex-bead phagosomes. The majority of the BBCs float at the 10%-25% interphase.

**Table S1. *M. marinum*-MCV early proteome**

**Table S2. *M. marinum*-MCV early proteome KEGG pathways**

**Table S3. Proteins differentially abundant during *M. marinum*-MCV establishment**

**Table S4. Proteins differentially abundant in M. marinum-MCV compared to non or less-manipulated MCVs**

**Table S5. MS thresholds calculations**

## Notes

### Competing Interest Statement

The authors have declared no competing interest.

### Summary of Updates

The proteomics result have been re-analysed using a slightly different pipeline and a novel representation for the GO term enrichment analyses. In addition, a more comprehensive validation of major hits has been added, using expression and localisation of GFP fusions during infection. The discussion/conclusion section has been expanded.

